# N1-methylpseudouridine mRNA modification enhances efficiency and specificity of gene overexpression by preventing Prkra-mediated global translation repression

**DOI:** 10.1101/2024.12.06.627293

**Authors:** Tong Lu, Aijun Chen, Sen Wang, Yizhuang Zhang, Boya Yang, Jiasheng Wang, Hailing Fang, Qianqian Gong, Ang Li, Xiangguo Liu, Pengcheng Ma, Bingyu Mao, De-Li Shi, Ming Shao

## Abstract

In vitro transcribed messenger RNA (IVT mRNA) has emerged as a pivotal tool in mRNA-based therapies and has been extensively employed in gene function studies and genetic tool applications. However, the IVT process generates double-stranded RNA (dsRNA) by-products that are recognized by dsRNA sensors, triggering innate immune responses. In this study, we comprehensively analyzed the detrimental effects of dsRNA by-products on early zebrafish embryos, revealing that these by-products induce cell necrosis and delay maternal-zygotic transition by reducing global translation efficiency via Prkra, a recently identified dsRNA sensor in pluripotent cells. Importantly, we demonstrate that N1-methylpseudouridine (m1φ) modification of IVT mRNAs effectively mitigates these adverse effects, as m1φ-modified dsRNAs exhibit significantly lower binding affinity to the Prkra dimer. Our findings underscore a previously overlooked challenge in the use of IVT mRNA in early embryos and offer a robust solution to enhance the fidelity of mRNA applications. Furthermore, we elucidate that m1φ modification minimizes the dsRNA-induced stress response in pluripotent cells through a distinct mechanism.

## Introduction

In vitro transcribed messenger RNA (IVT mRNA) is extensively utilized in gene gain-of-function studies and serves as a fundamental component in mRNA-based therapies, with mRNA vaccines against SARS-CoV-2 being the most notable example.^1,2^ IVT process accompanies the production of double-stranded RNA (dsRNA) by-products,^3,4^ which are recognized as virus-like non-self molecules by pattern-recognition receptors (PRRs). These include endosomal toll-like receptors such as TLR3, as well as cytosolic sensors like retinoic acid-inducible gene I (RIG-I) and melanoma differentiation-associated 5 (MDA5).^5–7^ Activation of these PRRs triggers signaling pathways that elevate the expression of interferons (IFNs) and pro-inflammatory cytokines. They initiate outside-in JAK/STAT cascades that upregulate numerous genes, including interferon-stimulated genes (ISGs) involved in antiviral responses.^8,9^ Furthermore, dsRNA can activate an integrated stress response (ISR) by binding to and activating protein kinase R (PKR), which inhibits global protein translation through the phosphorylation of the translation initiation factor eIF2α.^10,11^ Uridine modifications, such as pseudouridine (φ) and 2-thiouridine (S2U), effectively diminish the recognition of dsRNA by sensors including RIG-I, PKR and TLR3.^12–16^ These strategic alterations enhance the potential for vaccination while minimizing associated inflammatory responses. Notably, N1-methylpseudouridine (m1φ) modification of mRNAs elicits an even lower immune response than φ while supporting enhanced translation activity.^17^

These conclusions are primarily drawn from studies on differentiated cells. However, our recent findings reveal that in pluripotent stem cells, dsRNA is detected by the dimerized dsRNA-binding protein Prkra, circumventing the canonical dsRNA sensors and PKR-eIF2α pathway.^18^ Prkra is activated by dsRNA through a highly ordered binding mechanism. Prkra-dsRNA complex leads to a global translation inhibition by the sequestration of translation machinery components, a process facilitated by dimerized dsRNA binding domain 3s (dsRBD3s).^18^ The impact of this recently identified dsRNA-induced stress mechanism on the utilization of IVT mRNA in early embryos and stem cells remains unexplored. Additionally, it is still unclear whether and how m1φ mRNA modification affects the Prkra-mediated stress response in undifferentiated cells.

The use of unmodified IVT mRNA in developmental biology has a long-standing history. The first mRNA injection into *Xenopus* oocytes dates back to 1983,^19^ while for zebrafish, the first mRNA overexpression record was published in 1994.^20^ Since then, mRNA-based gene overexpression has become a standard approach for gene gain-of-function studies and rescuing gene loss-of-function phenotypes. More importantly, genetic tools for transgenesis and genome editing depend highly on IVT mRNAs encoding exogenous functional proteins like Tol2 transposase, Cas9, etc.^21–23^ However, because the side effects of IVT mRNA are often interpreted as the function of its encoded protein, the risk of introducing IVT mRNA in early embryos has been largely ignored for decades.

In this study, we examined the detrimental effects of dsRNA by-products generated during the IVT of mRNA intended for gene overexpression and genetic tool application. These dsRNA by-products often lead to extensive cell necrosis, delay of maternal-zygotic transition and the emergence of misleading phenotypes, which can hinder the functional rescue by wild-type mRNA and reduce the effectiveness of mRNA-based tools in genome editing and transgenesis. Importantly, we found that m1φ modification significantly mitigates the toxic effects of dsRNA in early embryos. Mechanistically, m1φ modification markedly decreases the binding of Prkra to dsRNA, thereby alleviating the activation of Prkra-induced global translation repression in the early embryos. Our findings provide a robust solution for achieving accurate gene overexpression in early zebrafish embryos, enhancing the fidelity and reliability of mRNA-based genetic tools in this model system. Additionally, we present evidence and the underlying mechanism that m1φ modification minimizes the dsRNA-induced stress response in pluripotent cells.

## Results

### dsRNAs in IVT mRNAs induce toxic effect on zebrafish early embryos

Injection of IVT mRNAs into zebrafish embryos frequently cause toxicity, as indicated by massive deaths and malformed embryos. Although early cleavage was not affected (Video S1), *cas9* mRNA-injected embryos (200 pg/embryo) exhibited general developmental delay at the onset of epiboly movement (Figure 1A and Video S2). More than one-fifth of injected embryos lysed during the gastrula stage, probably due to disrupted cortical F-actin and microtubule cytoskeletons in the cortical layer of the yolk cell (Figures 1B, B’, C, C’). Injected embryos that survived at 36 hours postfertilization (hpf) exhibited developmental defects, such as tail truncation, split axis, abnormality or absence of eyes, malformed brain structures, and cardiac edema (Figure 1D). We categorized the phenotypes as follows: normal (embryos exhibiting typical morphology without noticeable defects); mild (embryos displaying a slightly shortened body axis, with or without accompanying pericardial edema); moderate (embryos characterized by a significantly shortened body length, with or without ocular defects); severe (embryos with complete absence of the trunk and tail, as well as head deformities or a split body axis).

**Figure 1.**
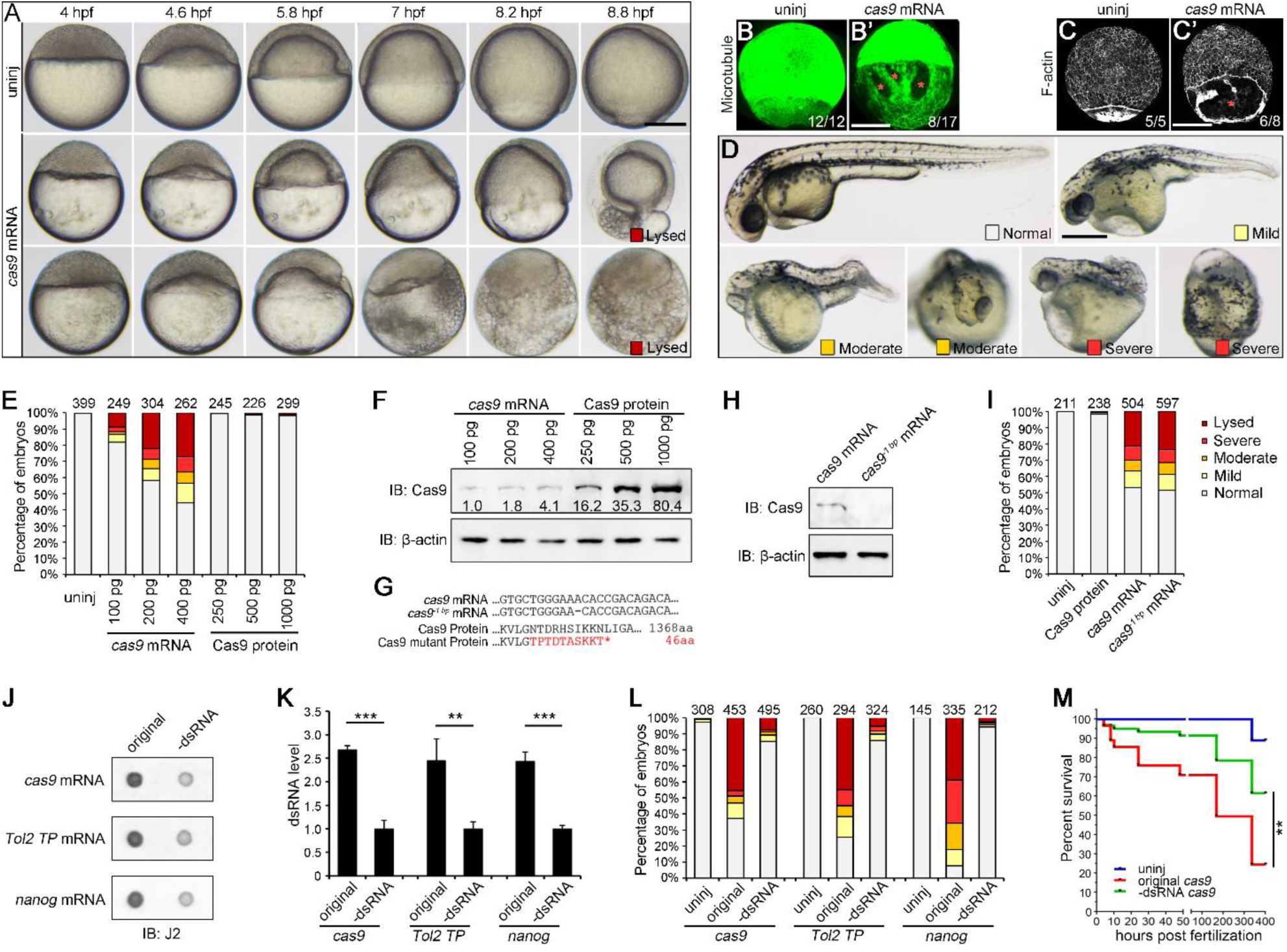
dsRNA in IVT mRNA triggers unspecific toxic effects in zebrafish embryos. (**A**) Time-lapse imaging of *cas9* IVT mRNA-injected embryos, highlighting yolk cell rupture during gastrulation. (**B**), (**B’**), (**C**), and (**C’**) α-tubulin antibody staining (B and B’), and phalloidin staining (C and C’). Asterisks mark the cytoskeleton-absent “holes” on the surface of the yolk cell. (**D**) Aberrant phenotypes of *cas9* mRNA-injected embryos at 36 hpf. (**E**) Phenotypic analysis of *cas9* mRNA or Cas9 protein-injected embryos. (**F**) Western blot analysis to determine Cas9 protein level in mRNA and protein-injected groups. (**G**) The nucleotide sequence of 1 bp deletion in *cas9* mRNA and the resultant amino acid sequence. (**H**) Mutant *cas9* mRNA failed to generate Cas9 protein. (**I**) Phenotypic analysis of wild-type and −1 bp mutant *cas9* mRNA injected embryos. (**J**) Dot blot analysis with J2 antibody of three IVT mRNAs with or without cellulose chromatography. (**K**) Quantification of dot blot from three biological replicates. (**L**) Phenotypic analysis after injecting 200 pg/embryo IVT mRNAs. (**M**) Effects of *cas9* mRNAs on survival of zebrafish embryos. Values are means ± SDs. **** in **M**, p<0.0001, Log-rank (Mantel-Cox) test. For others, ** p<0.01, *** p<0.001, Student’s *t*-test. Scale bars: 200 μm.

Interestingly, these aberrant phenotypes depended on the dose of *cas9* IVT mRNAs but were not related to the Cas9 protein level (Figure 1E). The Cas9 protein levels in the protein-injected group were 4 to 80 folds higher than those in the mRNA-injected group (Figure 1F). However, such high levels of Cas9 protein seldomly resulted in malformed embryos (Figure 1E). Thus, components in the IVT mRNAs per se should cause these toxic effects. Consistently, *cas9* mRNA with one base pair (bp) deletion in the open reading frame (ORF) did not produce functional Cas9 protein (Figures 1G, H) but had similar toxicity when 400 pg were injected into the embryo (Figure 1I). We also compared the toxicity of mRNAs encoding Tol2 TP or Nanog with their frame-shifted counterparts and found that they elicited comparable teratogenic effects on early embryos (Figures S1A-C).

IVT often generates dsRNA by-products via 3’-end extension and antisense strand transcription.^24–27^ In addition, the toxic phenotypes resulting from IVT mRNAs are analogous to the malformations induced by dsRNAs.^18,28^ Thus, we speculated that the dsRNA contaminants in mRNA might cause the toxic effect. A noticeable phenomenon is that the toxic effects varied between different batches of IVT mRNAs (Figure S1D). By dot blot analysis using a dsRNA-specific J2 antibody, we found that *cas9*, *Tol2-TP*, and *nanog* mRNAs with substantial toxicity showed 2.6 to 3.6 folds more dsRNA contamination (Figures S1E, F). Interestingly, IVT mRNAs with varying sequences exhibited differing levels of toxicity, which appears to be positively correlated with the levels of dsRNAs (Figures S1G-I). We also detected the presence of reverse-complementary RNA sequences derived from both highly and weakly toxic *cas9* mRNAs through RT-PCR analysis, further confirming the presence of dsRNAs in toxic IVT mRNAs (Figures S1J, K). qRT-PCR also revealed that weakly toxic IVT *cas9* mRNAs had significantly lower levels of reverse-complementary RNAs (Figure S1L). Next, we used cellulose chromatography to reduce dsRNA byproducts in IVT mRNAs,^29^ resulting in a 60%-80% decrease in dsRNA levels (Figures 1J, K, and S1M). As expected, injection of purified mRNAs (200 pg/embryo) significantly reduced toxic effects (Figure 1L) and the removal of dsRNAs also led to a well-improved survival rate (Figure 1M).

### dsRNAs elicit severe cell necrosis and delay the maternal-zygotic transition

The dsRNA-induced toxic phenotypes are possibly associated with cell death. As the toxicity of dsRNA is sequence independent, we next used a previously described 1 kb dsRNA from *cas9* open reading frame in the following analysis.^18^ In sections made from dsRNA-injected embryos at 8 hpf, we frequently observed enlarged and fragmented cells, reminiscent of cell necrosis (Figures 2A, B). Consistent with the frequent membrane rupture, F-actin lining the cell membrane was remarkably reduced in swollen cells after dsRNA injection (Figures 2C, D). To further confirm the presence of necrosis, we stained dsRNA-injected (50 pg/embryo) live embryos with propidium iodide (PI) and observed red fluorescent nuclei gradually emerging during development (Figures 2E, F, Video S3), suggesting the loss of membrane integrity, a hallmark for necrosis. However, these gradually emerged PI-stained nuclei never appeared in uninjected controls. In cell transplantation experiments, large swollen and broken cells were also present in gastrula embryos (Figures 2G, H). The process of cell membrane rupture was also captured in isolated single cells (Figure 2I, Video S4). We also compared the ultrastructure between dsRNA-stimulated and uninjected cells by transmission electron microscopy (TEM) at 8 hpf. Swollen cells caused by the injection of dsRNAs exhibited active mitophagy (Figure S2), suggesting a potential self-rescue response to mitigate the stress caused by dsRNA stimulation.

**Figure 2.**
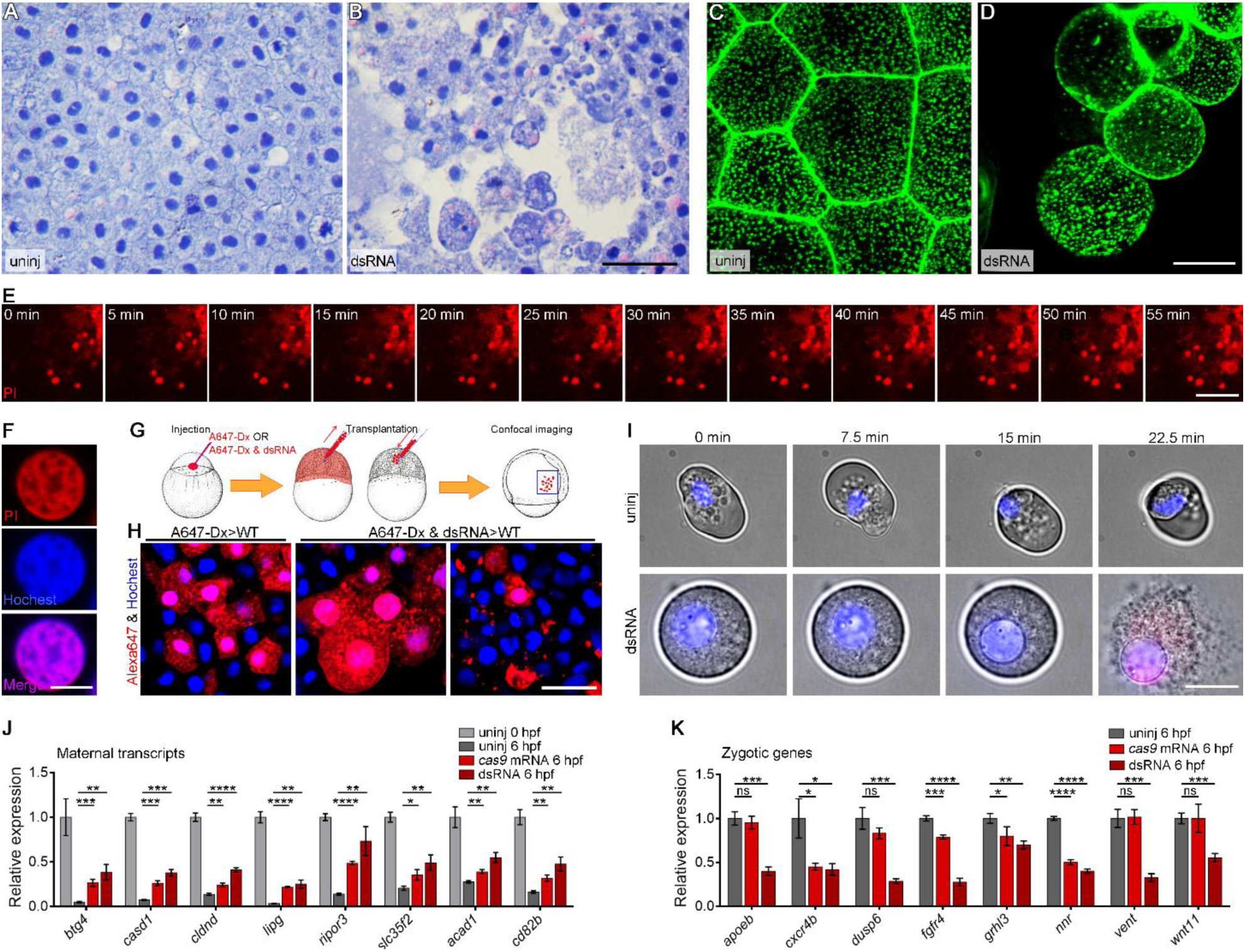
dsRNAs induce cell necrosis and delay MZT. (**A** and **B**) JB4 sections of uninjected and dsRNA-injected embryos at 8 hpf stained by HE. (**C** and **D**) Phalloidin staining to visualize cortical F-actin in uninjected (**C**) and dsRNA stimulated (**D**) cells at 8 hpf. (**E** and **F**) The emerging red dots are PI-stained nuclei in a time-lapse analysis of a dsRNA-injected embryo. (**G**) Procedure of labeling, transplantation and imaging. (**H**) dsRNA-injected cells exhibit swelling and fragmentation in chimera embryos. (**I**) Time-lapse analysis showing the rupture process of single isolated cells from embryos injected with dsRNA. (**J**) qRT-PCR analysis of eight maternal genes’ expression in uninjected, IVT mRNA injected (100 pg/embryo) and dsRNA injected (30 pg/embryo) embryos. (**K**) qRT-PCR analysis of the transcript level of eight zygotically expressed genes at 6 hpf. Values shown in columns are means ± SDs. ns, p > 0.05; * p<0.05; **, p < 0.01; ***, p < 0.001; ****, p < 0.0001, Student’s *t*-test. Scale bars: 50 μm for **A**, **B** and **H**; 20 μm for **C**, **D** and **I**; 100 μm for **E**, 10 μm for **F**.

Given the significant developmental delay observed following dsRNA injection, we investigated whether dsRNA interfered with maternal-zygotic transition (MZT), also called zygotic genome activation (ZGA) or mid-blastula transition (MBT), representing a crucial event during early development. In this critical period, a substantial proportion of maternal transcripts are typically degraded, while zygotic transcripts rapidly accumulate from the activated genome. To determine the effects of dsRNAs on these processes, we assessed the expression of eight maternal transcripts, *btg4*, *casd1*, *cldnd*, *lipg*, *ripor3*, *slc35f2*, *acad1* and *cd82b*, which typically undergo rapid degradation following MZT in uninjected embryos.^30^ qRT-PCR analysis revealed significantly reduced degradation of these transcripts after dsRNA stimulation, either from annealed dsRNAs or IVT *cas9* mRNAs (Figure 2J). In addition, we examined the expression of eight zygotic genes, including *apoeb*, *cxcr4b*, *dusp6*, *fgfr4*, *grhl3*, *nnr*, *vent*, and *wnt11*.^30^ Injection of annealed dsRNAs (30 pg/embryo) inhibited the transcriptional activation of all these genes at 6 hpf, and injection of dsRNA-positive IVT mRNAs (200 pg/embryo) also showed a similar trend, though to a lesser extent (Figure 2K). These findings suggest a notable delay in the MZT following dsRNA stimulation.

### Global translation inhibition mediated by Prkra is responsible for the toxic effects of IVT mRNA

Given that dsRNA stimulation can induce global translation inhibition,^18^ we sought to determine whether defects in protein synthesis are sufficient to elicit non-specific toxic phenotypes. For this purpose, we treated zebrafish embryos with the translation inhibitors cycloheximide (CHX) and puromycin, which inhibit translation elongation, for comparison with the effects of dsRNA injection. Our results indicated that the translation repression induced by these two compounds mimicked the effects of dsRNAs on overall phenotypes and necrosis when administered at titrated concentrations (Figures 3A-D).

**Figure 3.**
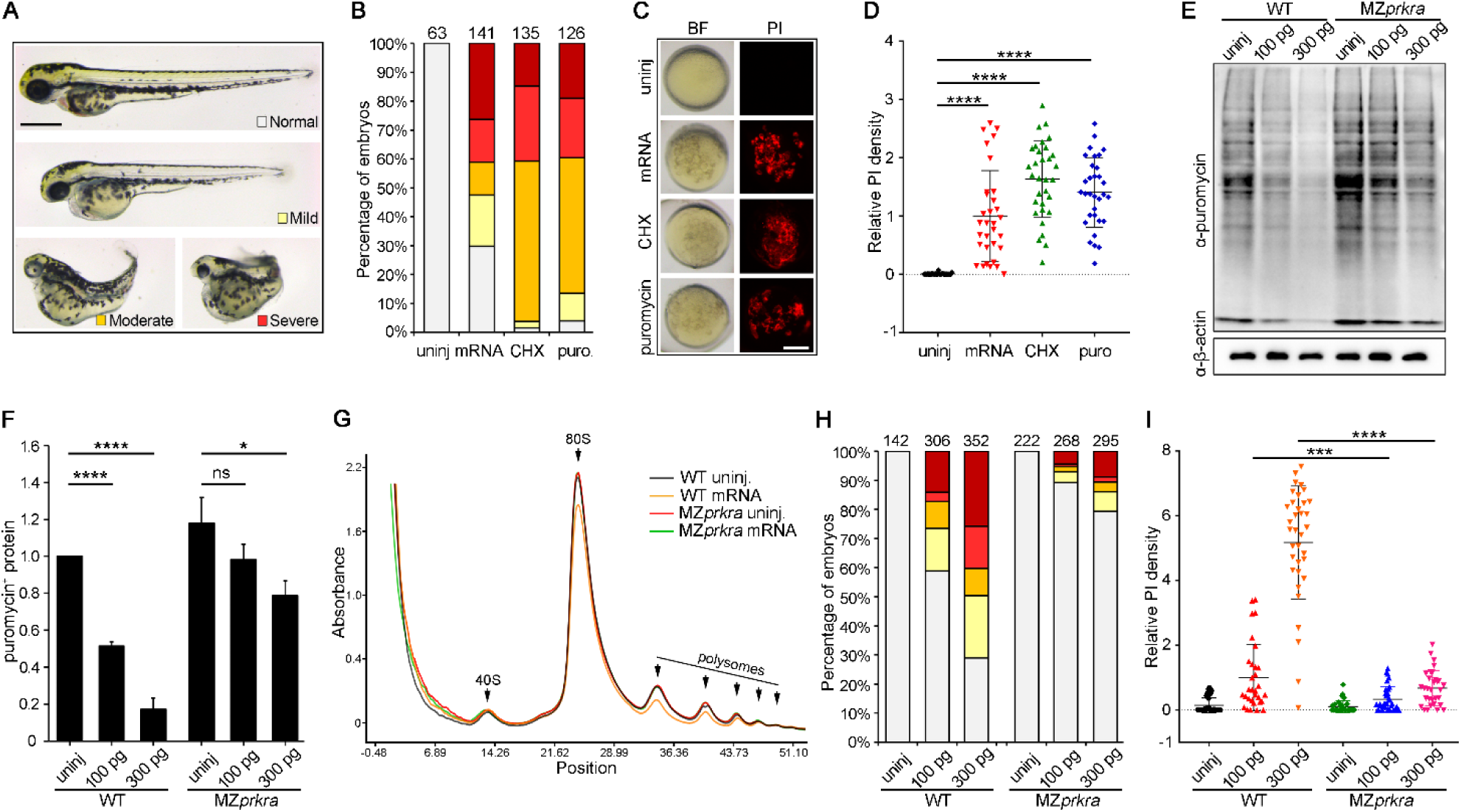
The toxic effects of IVT mRNA are dependent on the Prkra-mediated dsRNA sensing and global translation repression. (**A**) Representative phenotypes of embryos treated with translation inhibitors, CHX and puromycin. (**B**) Quantitative phenotypic analysis of embryos injected with dsRNA (30 pg/embryo) or treated with CHX (5 μg/mL) or puromycin (8 μg/mL). The numbers above each stacked column represent the total number of embryos analyzed. (**C**) dsRNA injection and treatment with CHX and puromycin cause cell necrosis in 10 hpf embryos, as revealed by PI staining. (**D**) Quantification of fluorescence intensity of PI in individual embryo. (**E**) Puromycin incorporation assay showing translation inhibitory effects of dsRNA in wild-type and MZ*prkra* embryos. (**F**) Quantification of incorporated puromycin levels. (**G**) Ribosome profile analysis via sucrose gradient ultracentrifugation. Notably, dsRNA treatment reduces the polysome fraction in wild-type embryos, but not in MZ*prkra* embryos. (**H**) Phenotypic analysis of dsRNA-injected wild-type and MZ*prkra* embryos. (**I**) Comparison of cell necrosis between wild-type and MZ*prkra* embryos following dsRNA injection. Values shown in columns are means ± SDs. ns, p > 0.05; * p<0.05; ***, p < 0.001; ****, p < 0.0001, Student’s *t*-test. Scale bars: 500 μm for **A**; 200 μm for **C**.

In early embryos and embryonic stem cells, the dsRNA sensor involved in regulating global translation efficiency is Prkra, rather than PKR.^18^ To investigate whether mRNA injection induces translation blockage in a Prkra-dependent manner, we examined the effects of IVT mRNAs in wild-type and MZ*prkra* embryos. The MZ*prkra* embryos were generated by crossing homozygous mutants carrying two complex mutations (Figures S3A, B). One mutant has −5 bp, −8 bp, and −14 bp deletions at the first three sgRNA target sites, with the initial −5 bp deletion disrupting the exon-intron junction (Figure S3B). The other mutant allele contains a large 235 bp deletion flanked by the second and third sgRNA target sequences, as well as −6 bp and −9 bp small deletions at the first and fourth sgRNA sites (Figure S3B). Both mutations prevent the production of a functional Prkra protein (Figure S3C) and lead to nonsense mediated decay (NMD) (Figure S3D). By injecting 100 and 300 pg *cas9* IVT mRNAs into wild-type and MZ*prkra* embryos, we found that unlike wild-type embryos, where approximately 50-80% translation inhibition was observed, translation inhibition was significantly alleviated in *prkra* mutants (Figures 3E, F). The ribosome profile analysis through ultracentrifugation using 5-hpf embryos revealed that the administration of 100 pg/embryo of *cas9* IVT mRNAs reduced the polysome fraction in wild-type embryos, but not in MZ*prkra* embryos (Figure 3G). This finding aligns with the observation that injecting dsRNA-positive IVT mRNAs into MZ*prkra* embryos produced minimal aberrant phenotypes and necrosis (Figures 3H, I). These results thus demonstrate that dsRNA-Prkra axis and the resultant translation inhibition is responsible for the toxic effects of IVT mRNAs.

### mRNA overexpression leads to experimental artifacts and compromise the efficiency of mRNA tools

The above results led us to realize that the dsRNA-induced stress response may impact the effectiveness and reliability of mRNA overexpression. Here, we present two examples to substantiate this assertion. The first is a co-injection experiment. We activated Nodal signaling by injecting dsRNA- removed *squint-myc* (*sqt-myc*) mRNA which encodes a myc-tagged Nodal ligand. This resulted in ubiquitous expression of Nodal target gene *goosecoid* (*gsc*). When we co-injected *nanog* mRNA along with *sqt-myc*, the ectopic induction of *gsc* was significantly inhibited (Figures 4A, B). This observation seemingly indicates a suppressive effect of Nanog protein on Nodal signaling. However, after removal of dsRNAs by cellulose chromatography, the purified *nanog* mRNA did not exhibit such an effect (Figure 4B), indicating that the above phenomenon only represents a stress effect from dsRNAs. In fact, the dsRNAs in *nanog* IVT mRNAs significantly reduced global translation efficiency and thus the production of Squint-myc protein, while the cellulose purified *nanog* mRNA did not (Figure S4A).

**Figure 4.**
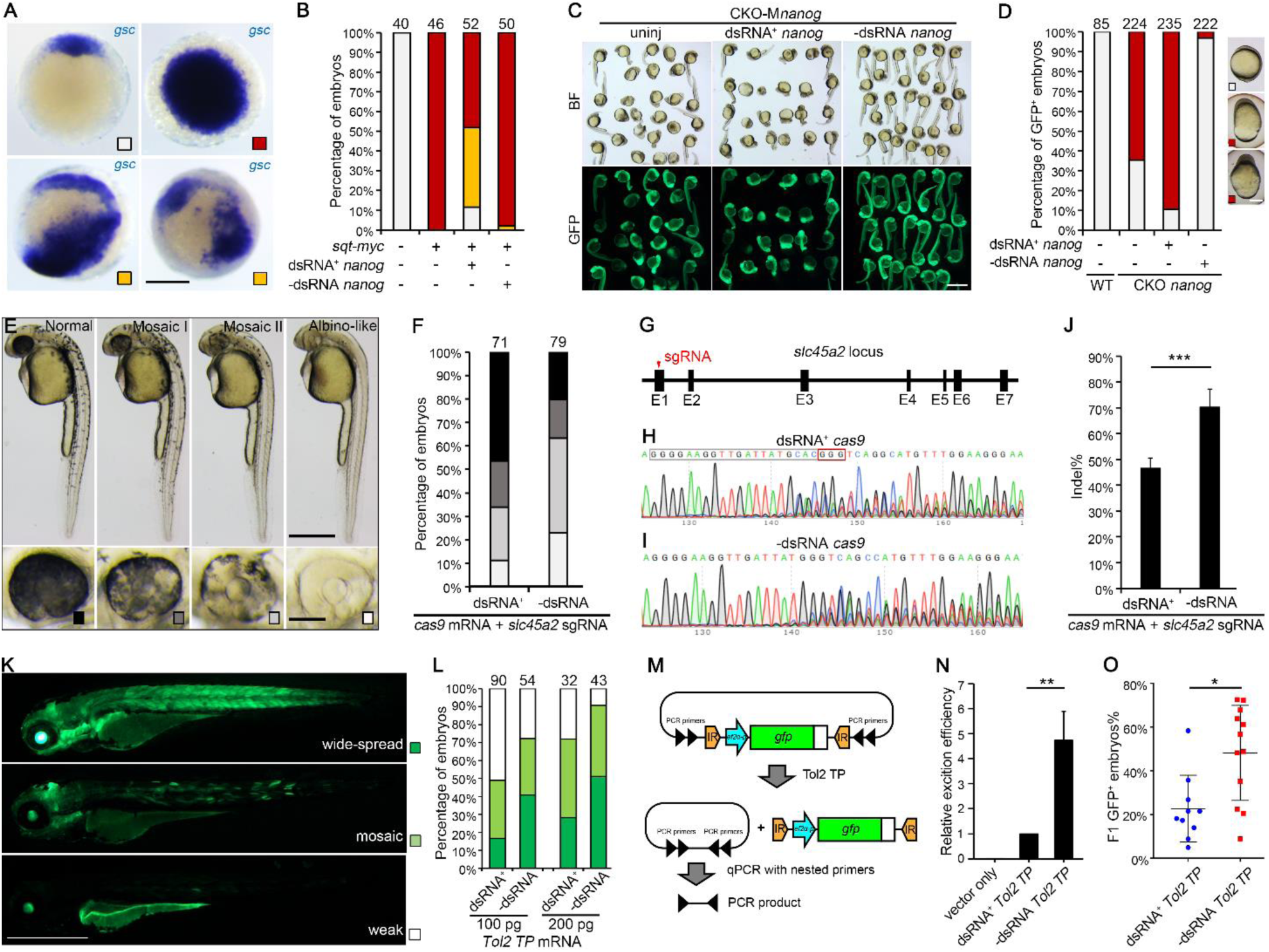
dsRNA by-products cause experimental artifacts and reduce the efficiency of genetic tools. (**A**) Typical ISH result of *goosecoid* in Nodal-overexpressed embryos at 6 hpf. (**B**) Phenotypic analysis shows that dsRNA-positive *nanog* IVT mRNA has an inhibiting effect of Nodal activation. (**C**) Phenotypes of CKO *nanog* maternal mutant, with or without injecting *nanog* IVT mRNAs. (**D**) dsRNA-removed *nanog* mRNA, but not the unpurified counterpart, can rescue the defective phenotype of its maternal mutant. (**E**) Representative phenotypes of 36-hpf *slc45a2* CRISPants. (**F**) Phenotypic analysis of *slc45a2* CRISPants. (**G**) Structure and sgRNA target site of *slc45a2* locus. (**H** and **I**) Sequencing chromatograph after editing by coinjecting *slc45a2* sgRNA and unpurified (H) or dsRNA-removed (I) *cas9* IVT mRNA. (**J**) Quantification of editing efficiency by Tide algorithm. (**K**) Representative fluorescence patterns of 3 dpf transgenic F0 larva. (**L**) Quantification of fluorescence pattern after injecting unpurified and dsRNA-removed *Tol2 TP* mRNA. (**M**) The experimental design of transposon excision assay. (**N**) dsRNA removal significantly increased excision efficiency. (**O**) The percentage of GFP-positive offspring significantly increases after injecting dsRNA-removed *Tol2 TP* mRNA. Values shown in columns are means ± SDs. * p<0.05; **, p < 0.01; ***, p < 0.001, Student’s *t*-test. Scale bars: 200 μm for **A** and **D**; 500 μm for upper row in **E**, 100 μm for the lower row; 1 mm for **C** and **K**.

The adverse effect of dsRNA contaminants was also shown in a rescue experiment. We generated *nanog* maternal mutant (M*nanog*) embryos via an oocyte-specific conditional knockout approach.^31^ One founder female fish produced 65% M*nanog* mutants, which displayed severe epiboly defect and a dorsalized-like phenotype (Figure 4C). The dsRNA-removed *nanog* mRNA can efficiently rescue these mutant phenotypes. Instead, injecting the dsRNA-positive *nanog* mRNA failed to rescue but exaggerated the lethal phenotypes (Figures 4C, D). These results thus demonstrate a considerable risk in using dsRNA-positive IVT mRNAs for gene functional studies in zebrafish embryos.

We also suspected that this stress response likely influences the effectiveness of mRNA tools. To test this, we first compared genome editing efficiencies by injecting the original and dsRNA-removed *cas9* mRNAs. By targeting *slc45a2*, which encodes a transporter protein that mediates melanin synthesis, we observe a remarkable increase in embryos with pigment loss phenotype when purified *cas9* mRNA was used (Figures 4E, F). Consistently, we observed a 1.5 folds higher mutation rate after injecting dsRNA-removed *cas9* mRNA than the unpurified control (Figures 4G-J). Similar improvement was observed when targeting the last intron of the *rbm24a* gene (Figures S4B-E). In accordant with the increased efficiency, purified *cas9* mRNAs produced three folds more Cas9 protein than the unpurified control (Figures S4F, G).

To test the effect of dsRNA by-products on the efficiency of transgenesis, we performed a proof-of-principle test using the pT2AL200R150G plasmid containing a ubiquitously expressed GFP reporter.^32^ When this vector was coinjected with *Tol2 transposase* (*Tol2 TP*) mRNAs, we observed an remarkably improvement of transgenic GFP expression at 3 dpf after coinjecting dsRNA-removed *Tol2 TP* mRNA (Figures 4K, L). Consistently, by transient embryonic excision assay,^32^ we detected a four-fold more increase in excision events after injecting dsRNA-purified *Tol2 TP* mRNA (Figures 4M, N). Also, the proportion of GFP-positive F1 offspring increased by almost one fold when using dsRNA-removed *Tol2 TP* mRNA (Figure 4O).

### m1φ modification mitigates the toxic side-effects of IVT mRNAs

The removal of dsRNAs present a viable approach to mitigate the toxicity associated with IVT mRNAs; however, the efficacy and reproducibility of the cellulose chromatography method for dsRNA removal are often constrained. Moreover, the yield of mRNAs is frequently below 50% following the complex and cumbersome cellulose purification process (see Materials and Methods), rendering this method less than optimal. Injecting unpurified IVT mRNAs into MZ*prkra* zebrafish can effectively mitigate or prevent toxicity; however, mutations in *prkra* may adversely impact zebrafish health and introduce potential risks when interpreting experimental results derived from this genetic background. Consequently, we sought to address the issue of gene overexpression by developing other strategies, with mRNA modification emerging as a critical option. Given that m1φ modification has been demonstrated to inhibit the activation of innate immune responses in differentiated cells,^17^ we hypothesized that this modification could similarly circumvent stress responses in undifferentiated cells and early embryos.

Therefore, we first compared the translation-inhibiting effects of m1φ-modified dsRNAs with its UTP analog and found that m1φ modification significantly diminished translation inhibition in wild-type embryos, as revealed by puromycin incorporation experiment (Figures 5A, B). As expected, neither UTP nor m1φ dsRNA could inhibit global translation in MZ*prkra* background (Figures 5A, B), which is consistent with the phenotypic and cell necrosis assay that m1φ dsRNAs failed to induce evident malformation and cell necrosis in either wild-type or MZ*prkra* background (Figures 5C, D). These results suggest that m1φ modification may enable evasion from the surveillance of dsRNA by its sensor, Prkra, in the early embryo.

**Figure 5.**
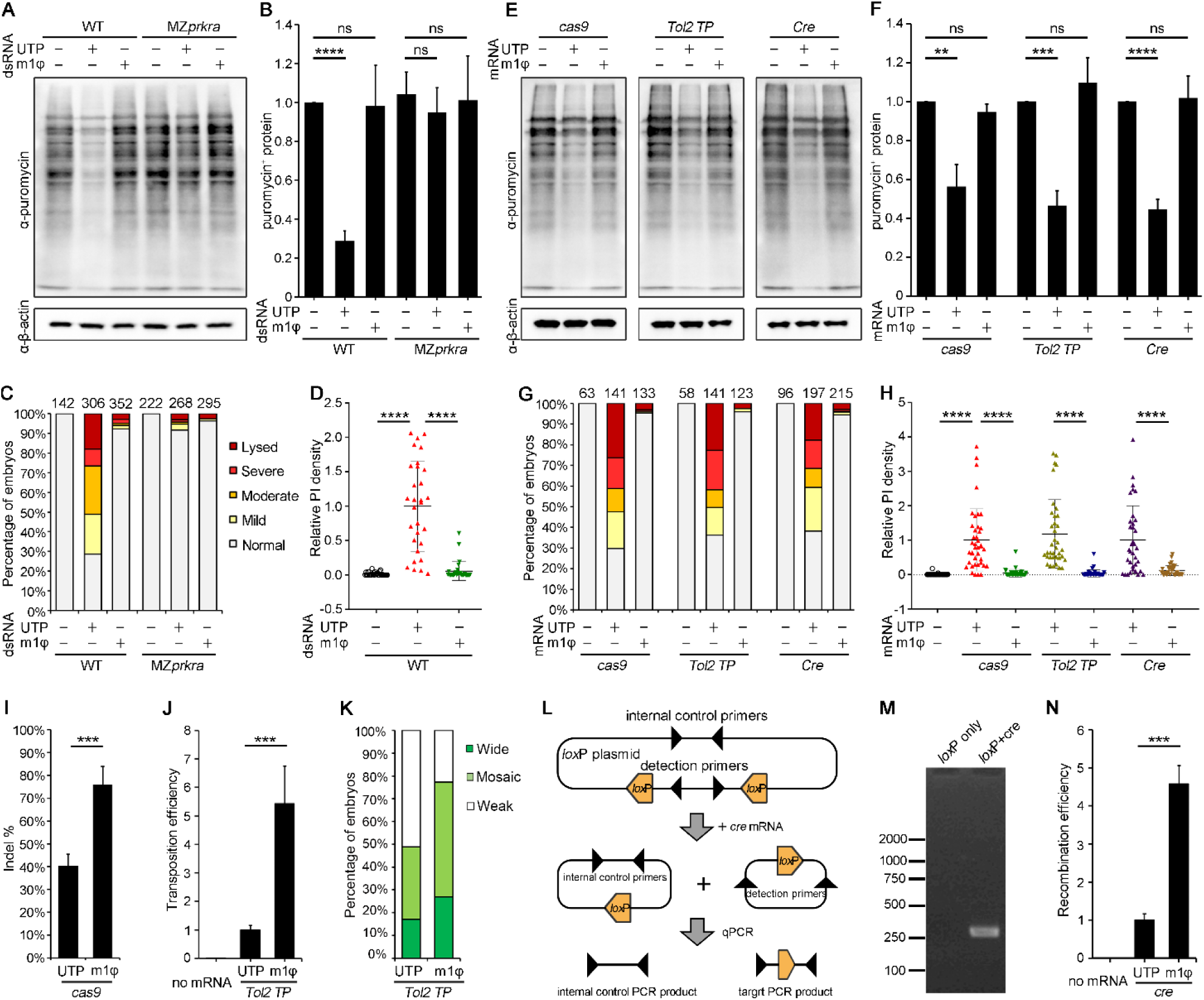
m1φ modification abolishes the dsRNA stress response and enhances the efficiency of mRNA-based genetic tools. (**A**) Puromycin incorporation assay demonstrating the loss of translation inhibition by m1φ-modified dsRNA in both wild-type and MZ*prkra* backgrounds. (**B**) Quantification of puromycin incorporation into newly synthesized proteins. (**C**) Phenotypic analysis of wild-type and MZ*prkra* embryos injected with m1φ or UTP dsRNA. (**D**) PI staining analysis showing significantly reduced cell necrosis induced by m1φ-modified dsRNAs. (**E**) Comparison of the translation-repressive effects of UTP and m1φ-modified *cas9*, *Tol2 TP*, and *cre* mRNAs injected at 200 pg/embryo. (**F**) Quantification of puromycin incorporation from panel **E**. (**G**) Phenotypic analysis of embryos injected with m1φ and unmodified IVT mRNAs. (**H**) Significantly reduced cell necrosis in embryos injected with m1φ-modified IVT mRNAs. (**I**) Elevated genome editing efficiency observed when injecting m1φ-modified *cas9* mRNA together with sgRNA targeting *slc45a2*. (**J**) Improved excision efficiency associated with the use of m1φ-modified *Tol2 TP* mRNA. (**K**) Increased transgenic GFP expression following the injection of m1φ *Tol2 TP* mRNA. (**L**) Experimental design for detecting recombination events using Cre. (**M**) PCR products from detection primers appear only when *cre* mRNA is co-injected. (**N**) Quantification of the recombination rate by qPCR. Values are expressed as means ± SD. ns, p > 0.05; **, p < 0.01; ***, p < 0.001; ****, p < 0.0001, Student’s *t*-test.

As such, we anticipated that this modification can eliminate the side effects of IVT mRNAs. We therefore synthesized *cas9*, *Tol2 TP*, and *cre* mRNAs using either UTP or m1φ. When injected at 300 pg/embryo, unmodified mRNAs significantly suppressed protein synthesis to approximately 50%, while m1φ-modified mRNAs at comparable levels exhibited no such an effect (Figures 5E, F). Consistently, high doses of UTP mRNAs induced significant toxic effects and a high incidence of lethal malformations. In stark contrast, the introduction of m1φ modification resulted in nearly normal development (Figure 5G). This observation aligns with the phenomenon whereby only UTP mRNAs triggered pronounced cell necrosis, while m1φ-modified mRNAs did not exhibit such effects (Figure 5H).

Hence, it can be expected that m1φ modification may help to increase the efficiency of mRNA-based genetic tools. We first assessed the genome editing efficiency by coinjecting UTP or m1φ- modified *cas9* mRNAs with sgRNA targeting the *slc45a2* gene, finding that the efficiency increased by about one fold with m1φ-modified *cas9* mRNA (Figure 5I). Similarly, when comparing the excision efficiency of Tol2 transposase translated from UTP and m1φ-modified mRNAs, we observed a more than five folds increase with m1φ-modified mRNAs (Figure 5J), which was associated with an improved transgenic GFP expression in 3 dpf embryos (Figure 5K). The *cre* mRNA represents another frequently used tool for removing *lox*P-flanked sequences. By co-injecting a *lox*P-containing plasmid with *cre* mRNA, with or without m1φ modification, we found a more than four folds increase in cutting efficiency by Cre protein translated from the modified *cre* mRNAs (Figures 5L-N). These results indicate that the use of m1φ-modified IVT mRNAs can effectively mitigate the toxic side effects, while also achieving a much more robust activity in the application of genetic tools.

### m1φ-modified dsRNA exhibits reduced binding affinity to the dsRNA sensor Prkra to mitigate translation inhibition

The above findings indicate that m1φ modification can effectively evade translation repression mediated by Prkra. To elucidate the underlying molecular mechanism, we hypothesized that m1φ modification might influence the binding affinity of the Prkra dimer to dsRNA. To investigate this, we synthesized fluorophore-labeled 85 bp AU-rich dsRNAs derived from sequences of the HIV genome (see Table S1), either with or without m1φ modification. We then conducted electrophoretic mobility shift assays (EMSA) to evaluate their binding characteristics to the Prkra dimer. Remarkably, the m1φ modification significantly weakened the binding affinity with the Prkra dimer (Figure 6A). Additionally, the typical laddering pattern resulted from the interaction of Prkra dimer with dsRNA was disrupted at high molecular weight level (Figure 6A), indicating that the highly ordered binding of the Prkra dimer is compromised following m1φ modification of dsRNA. Analysis of the binding curves revealed that the dissociation constant (Kd) for the m1φ-modified dsRNA was reduced to about 1/6 of that for unmodified UTP dsRNA (Figure 6B). However, the Hill coefficient (h) and maximum specific binding (Bmax) remained largely unchanged post-modification (1.005 vs 1.023), with the h value approximating 2 for both UTP dsRNA and m1φ-modified mRNA (2.125 vs 2.087). This suggests that binding of the Prkra dimer promotes further dimer binding to dsRNA, and this feature was not affected by m1φ modification.

**Figure 6.**
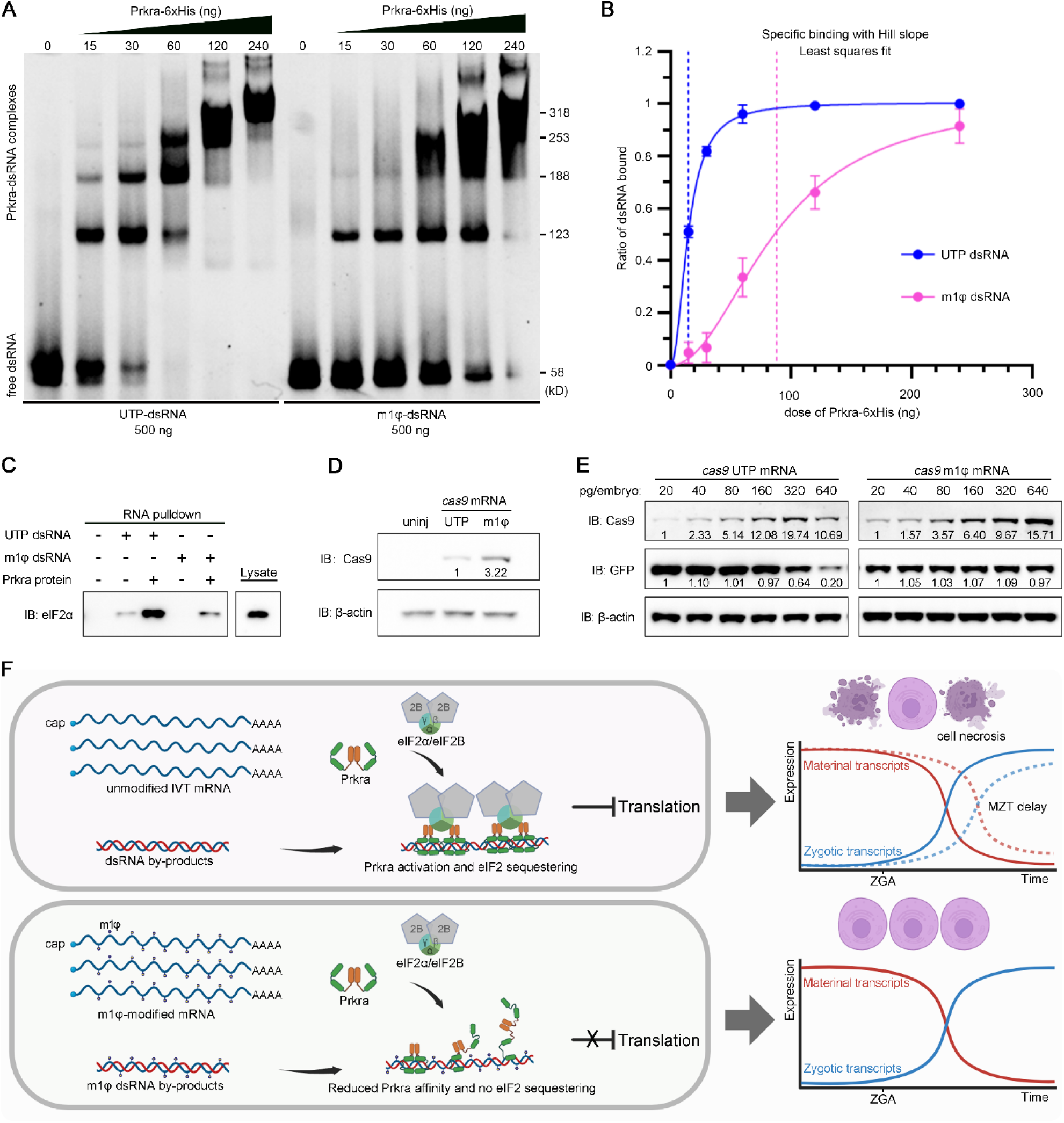
Reduced affinity of m1φ dsRNAs for Prkra dimer and enhanced translation of m1φ IVT mRNAs in early zebrafish embryos. (**A**) EMSA analysis of the binding patterns of UTP dsRNA and m1φ dsRNA to recombinant Prkra protein. (**B**) Binding curves showing the interaction of Prkra with UTP dsRNA and m1φ dsRNA. The vertical dashed lines indicate the dissociation constants for UTP and m1φ dsRNAs with the Prkra protein. (**C**) RNA pulldown assay demonstrates a significantly weakened interaction between m1φ dsRNA and eIF2α, both in the presence and absence of exogenous Prkra protein. (**D**) Western blot analysis comparing Cas9 protein production from UTP and m1φ *cas9* mRNAs. (**E**) Dose-dependent effects of UTP *cas9* mRNA and m1φ *cas9* mRNA on the translation of *gfp* reporter mRNA and their own translation. (**F**) Working model of the side-effects of IVT mRNA and the mechanism by which m1φ-derived dsRNA by-products compromise the stress response. In early embryos, dsRNA by-products are recognized by the Prkra dimer, which triggers global translation repression, ultimately leading to cell necrosis and a delay in MZT. In contrast, m1φ-derived dsRNA by-products fail to activate the Prkra sensor, thereby impairing the stress response.

We subsequently assessed the efficiency of dsRNA in sequestering the eIF2 complex through a dsRNA pull-down assay.^18^ As anticipated, m1φ modification significantly diminished the sequestration of eIF2α by dsRNA (Figure 6C). The addition of recombinant Prkra protein to the pull-down assay significantly enhanced the capture of eIF2α by dsRNA.^18^ Notably, the amount of eIF2α pulled down by m1φ-modified dsRNA is significantly reduced compared to that captured by the unmodified control in the presence of recombinant Prkra protein (Figure 6C). These results suggest that the modified dsRNA by-products may be not able to affect translation efficiency. Thus, the m1φ-modified IVT mRNAs likely display improved translation rates. Indeed, when injecting at 500 pg/embryo, the Cas9 protein translated from m1φ-modified mRNA was three times higher than that from the unmodified control, as revealed by western blot analysis (Figure 6D). To further evaluate the effect of mRNA dosage on the production of the protein of interest, we injected unmodified and m1φ-modified *cas9* IVT mRNAs at increasing doses, alongside with *gfp* mRNA at a constant dose of 30 pg/embryo. We observed that as the dose of UTP *cas9* mRNA increased, the amount of Cas9 protein initially rose but then declined when *cas9* mRNA was injected at extremely high doses (Figure 6E, left panel). In line with this trend, the translation efficiency of the co-injected *gfp* mRNA began to decrease to 64% when UTP *cas9* mRNA was injected at 320 ng/embryo, further dropping dramatically to 20% at a dose of 640 ng/embryo. Remarkably, for m1φ-modified *cas9* mRNA, the level of translated Cas9 protein consistently increased with the amount of mRNA injected, even at extremely high doses. Simultaneously, the translation rate of co-injected *gfp* mRNA remained unaffected (Figure 6E, right panel). These findings illustrate that m1φ-modified dsRNA byproducts generated during IVT exhibit significantly reduced binding affinity to Prkra, leading to decreased activation of Prkra-mediated sequestration of translation initiation factors (Figure 6F). We can thus conclude that m1φ-modified mRNAs are more suitable for applications in mRNA-based gene overexpression in zebrafish early embryos.

## Discussion

Our findings underscore the critical impact of dsRNA by-products generated during the IVT of mRNA, which induce global translation inhibition and misleading phenotypes in early zebrafish embryos. This study builds on our recent work, wherein we identified the dsRNA-binding protein Prkra as a mediator of translation repression, bypassing the canonical PKR-eIF2α pathway.^18^ Here, we deliver a thorough analysis of the detrimental effects of dsRNA by-products on gene overexpression studies and the application of genetic tools such as genome editing and transgenesis in the zebrafish model system, highlighting that dsRNA by-products induce cell necrosis and MZT delay (Figure 6F). Furthermore, we provided compelling evidence that m1φ modification of dsRNAs significantly reduces the binding and activation of Prkra. This discovery offers a robust solution to mitigate the adverse effects of dsRNA by-products, addressing a long-standing technical challenge in the early embryo.

In differentiated cells, achieving a complete absence of immunogenicity for mRNA requires both the modification of the RNA and the removal of dsRNA by-products.^33^ Modification of uridine is effective in reducing the affinity of dsRNAs to their sensors, such as RIG-I, PKR, and TLR3, as well as diminishing the affinity of single-stranded RNA (ssRNA) to TLR7 and TLR8.^12–16,34–37^ This reduction in affinity leads to decreased activation of downstream interferons and pro-inflammatory cytokines. However, m1φ- modified dsRNA can still partially activate the interferon response via RIG-I, mimicking the action of RIG-I agonist 5’ppp-hpRNA.^33^ Therefore, the elimination of any dsRNA contaminants is still crucial for the effective application of mRNAs in differentiated cells.

Several strategies have emerged to minimize or eliminate dsRNA byproducts in IVT mRNAs. One promising approach involves the engineering of T7 RNA polymerase to reduce dsRNA formation while maintaining high yields and purity of mRNAs. The double mutant T7 RNA polymerase (G47A + 884G) developed by Moderna exemplifies this strategy.^3^ Additionally, the covalent immobilization of T7 RNA polymerase and templates onto magnetic beads during IVT reactions has shown to effectively decrease dsRNA by-product levels.^38^ Further optimization of the IVT reaction conditions can also play a critical role. For instance, limiting UTP concentrations has been found to lower dsRNA levels, while restricting ATP concentrations paradoxically increases dsRNA formation.^39^ Moreover, the presence of loose mRNA secondary structures and an excess of unpaired uridine bases can lead to increased dsRNA production during IVT.^27^ This indicates that rNTP concentrations and sequence design are vital considerations for minimizing dsRNA contamination. This aligns with our observations that dsRNA production is influenced by the sequences of the templates and may also be related to the concentration of the template (Figures S1D-I). After the IVT process, several purification methods, such as high-performance liquid chromatography (HPLC) and cellulose chromatography, can be employed to remove dsRNA byproducts.^4,29^ Treatment with RNase III to cut dsRNA into small fragments has also showed an effect to reduce the immune-stimulatory response of mRNAs.^40^

In contrast to differentiated cells, it appears that m1φ modification alone is sufficient to circumvent the stress response in early zebrafish embryos, thus negating the need for further dsRNA removal. This phenomenon may be attributed to the low expression levels of RIG-I in early embryos, rendering it largely ineffective. This is consistent with the observation that dsRNA has a very low capacity to activate interferon gene expression in undifferentiated cells, a characteristic highly conserved across species, including zebrafish and mouse early embryos, mouse embryonic stem cells (mESCs), mouse teratocarcinoma cell lines, as well as human embryonic stem cells (hESCs) and induced pluripotent stem cells (hiPSCs).^18,41,42^ In contrast to RIG-I, the stem cell-specific dsRNA sensor Prkra does not recognize the triphosphate structure at the 5’ end of dsRNAs; instead, it is only able to recognize the conformation of dsRNA helices through its dsRBDs,^18^ thereby making m1φ modification of dsRNA sufficient to evade its detection.

Indeed, our findings demonstrate a significant reduction in the binding affinity between m1φ- modified dsRNA and Prkra, decreasing to approximately one-sixth that of UTP dsRNA. Additionally, the m1φ modification disrupts the laddering pattern observed in EMSA assays, indicating an alteration in the ordered arrangement of Prkra dimers on modified dsRNAs. As a result, m1φ-modified dsRNA fails to effectively capture the eIF2 complex in the presence of Prkra, underscoring its diminished interaction with key translational components. Consequently, the injection of m1φ-modified IVT mRNA allows for the administration of high doses without eliciting stress responses, even in contexts where dsRNA contamination is a concern. It was recently reported that the presence of m1φ may introduce a risk of translational frameshifting, resulting in truncated or misfolded protein products.^43^ Nevertheless, given the relatively low proportion of erroneous translations, the impact on the overall efficacy of gene overexpression may be minimal. Actually, our western blot analysis did not reveal the presence of any low molecular weight bands associated with translational frameshifting in m1φ-modified *cas9* mRNAs. This demonstrates an overall fidelity of translation in this case.

In summary, dsRNA by-products in IVT-synthesized mRNAs induce toxicity in early zebrafish embryos by activating Prkra dimer-mediated translation repression. The m1φ modification disrupts the recognition of dsRNA by the Prkra dimer, thereby alleviating the stress response. Given that the function of Prkra is conserved across species, it is conceivable that m1φ-modified mRNAs in mammalian stem cells can similarly evade this distinct innate immune surveillance, potentially facilitating more specific and robust gene overexpression.

## Supporting information

Raw data for each Figure

Uncropped gels and blots

## Data availability

Any information and requests for resources and reagents should be directed to and will be fulfilled by the corresponding author, Ming Shao.

## Supplementary Data statement

Figures S1-S4. Tables S1 and S2. Video S1-S4.

Data S1. (separate file) Raw data visualized in each Figure.

Data S2. (separate file) Uncropped blots presented in this paper.

## Acknowledgements

We thank J.L.S and H.J.D from the Core Facility and Service Platform, School of Life Sciences, H.Y.Y and X.M.Z from SKLMT for assistance with confocal imaging, histological sectioning and ribosome profiling, and Z.X.D for fish rearing.

## Supplementary Figures

**Figure S1.**
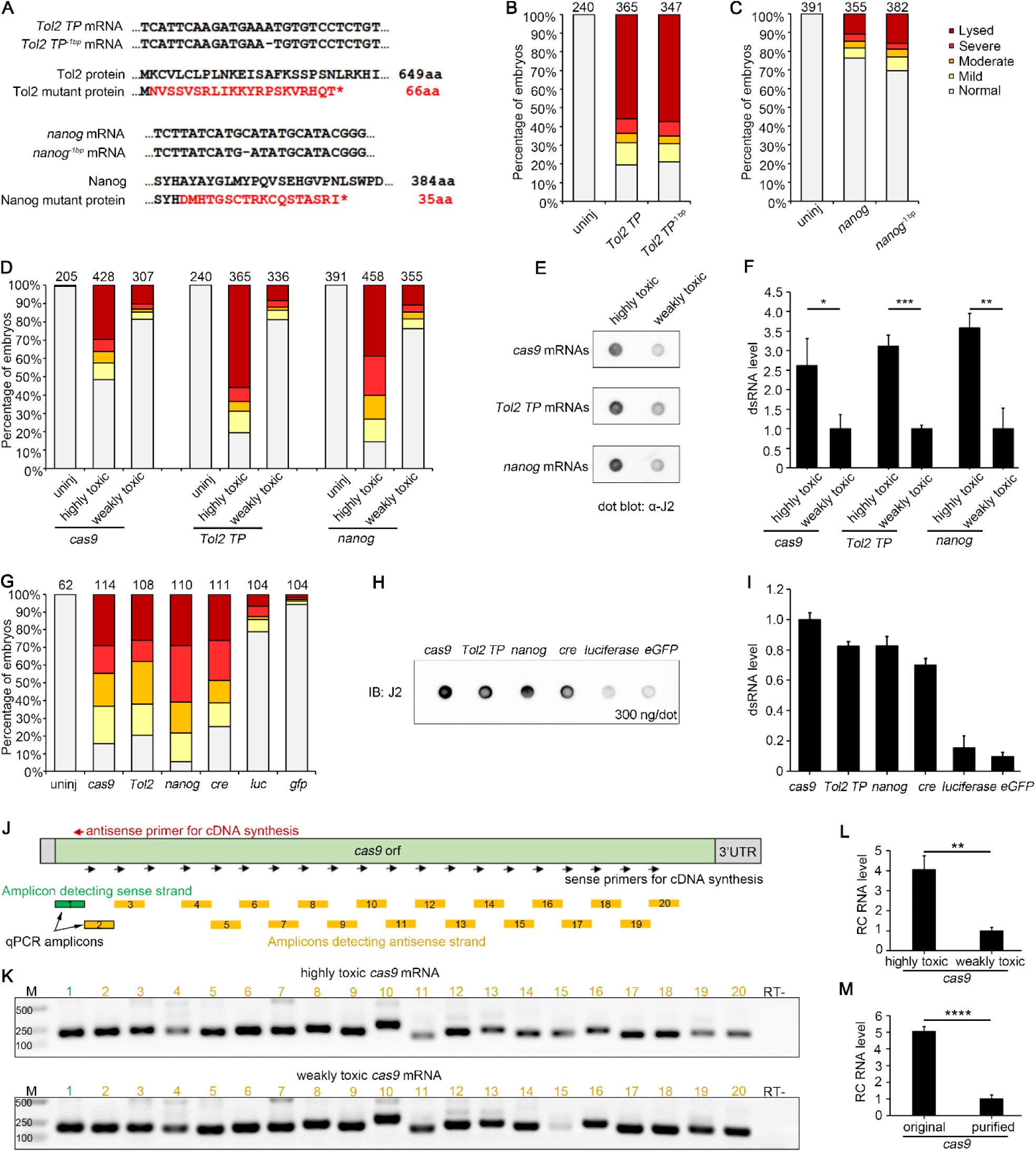
dsRNA positively correlates with the toxic effects of IVT mRNAs. (**A**) A 1 bp deletion in the *Tol2 TP* and *nanog* coding regions, along with the corresponding truncated proteins. (**B** and **C**) The toxicity of frameshift IVT mRNAs is comparable to that of wild-type controls. (**D**) Different batches of mRNA exhibit varying toxic effects on the embryos. (**E**) Dot blot analysis using J2 antibody reveals that highly toxic mRNA contains more dsRNA contaminants. (**F**) Quantification of dsRNA in highly and weakly toxic IVT mRNAs. (**G**) Phenotypic analysis demonstrates the differential toxic effects of IVT mRNAs with varying sequences. (**H**) Representative J2 antibody dot blot showing different levels of dsRNA in various IVT mRNAs. (**I**) Quantification of the dot blot shown in panel H. (**J**) A schematic illustrating the RT-PCR strategy for detecting antisense RNAs in *cas9* IVT mRNA. (**K**) RT-PCR results indicate the presence of antisense RNA in all examined regions of *cas9*. (**L**) The dsRNA level is significantly lower in weakly toxic *cas9* IVT mRNA. (**M**) The dsRNA level is significantly reduced following cellulose chromatography purification. The numbers at the top indicate the total number of embryos analyzed. Values are expressed as means ± SD. * p<0.05, ** p<0.01, *** p<0.001, **** p<0.0001, Student’s *t*-test.

**Figure S2.**
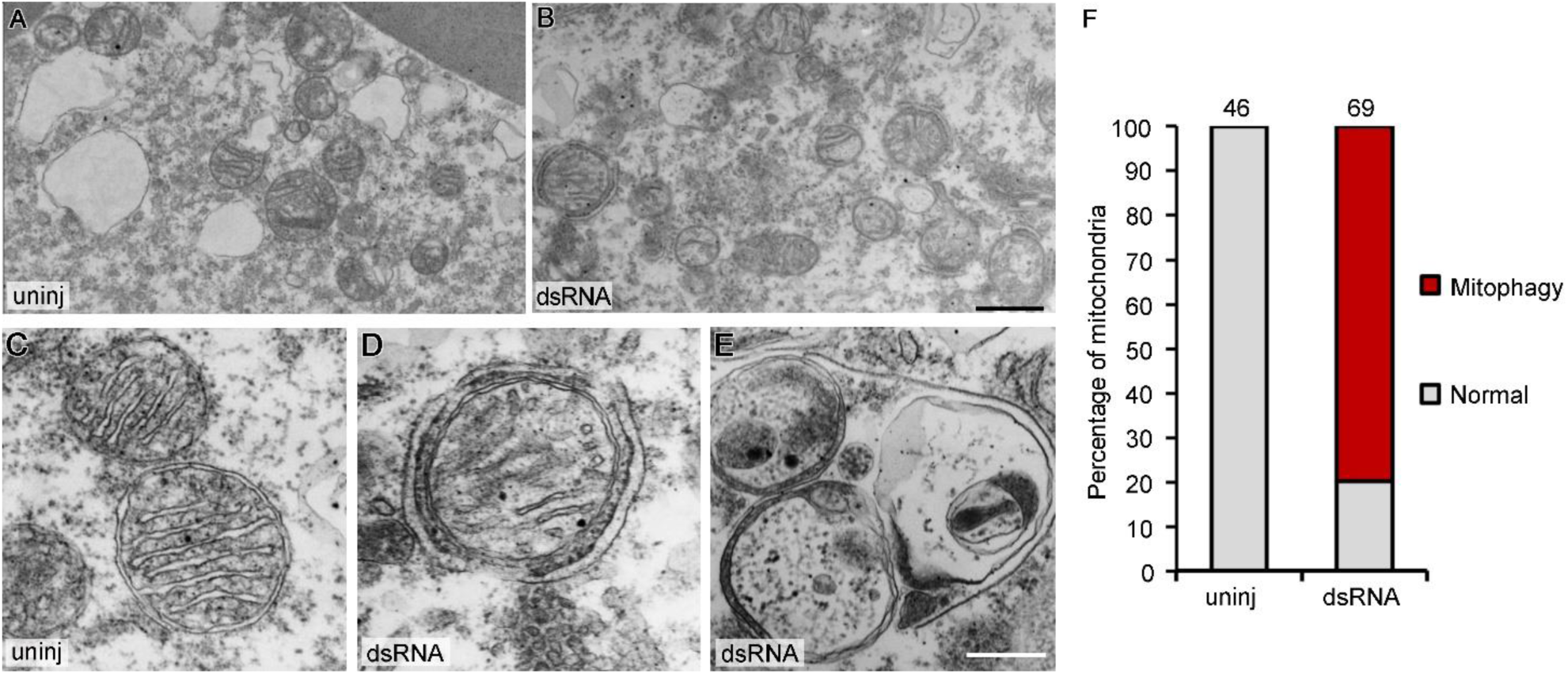
dsRNA stimulation induces mitophagy in early embryos. (**A**) Low magnification of the cytoplasmic region in a cell from an uninjected 8-hpf embryo. (**B**) Low magnification of the cytoplasmic region in a cell from a dsRNA-stimulated 8-hpf embryo. Notably, the majority of mitochondria are encircled by phagophores. (**C**-**E**) High magnification images of mitochondria in uninjected embryos and dsRNA-stimulated embryos. Panel D illustrates a mitochondrion surrounded by phagophores, while Panel E depicts degraded mitochondria within lysosomes. (**F**) Stacked columns demonstrating increased mitophagy in dsRNA-injected embryos. Numbers on top designate counted mitochondria. Scale bar: 500 nm for **A** and **B**; 200 nm for **D**.

**Figure S3.**
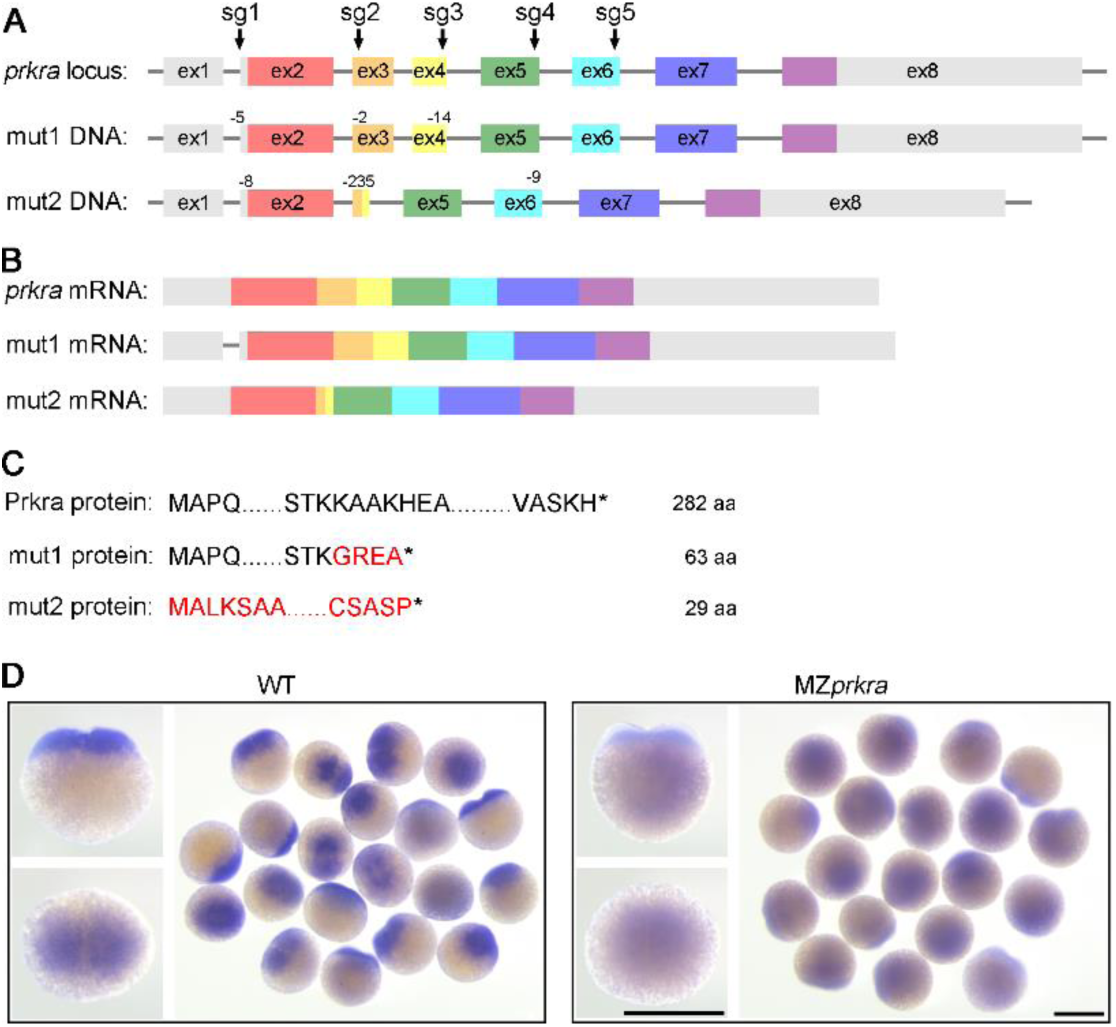
Null mutations in MZ*prkra* embryos. (**A**) Genome structure of wild-type and edited *prkra* loci. (**B**) Wild-type and mutant *prkra* mRNA generated by indicated alleles. (**C**) Predicted amino acid sequences of wild-type and mutant *prkra* mRNAs. (**D**) WISH experiments show that *prkra* mRNAs in MZ*prkra* mutant embryos undergo non-sense mediated decay. Scale bars: 500 μm.

**Figure S4.**
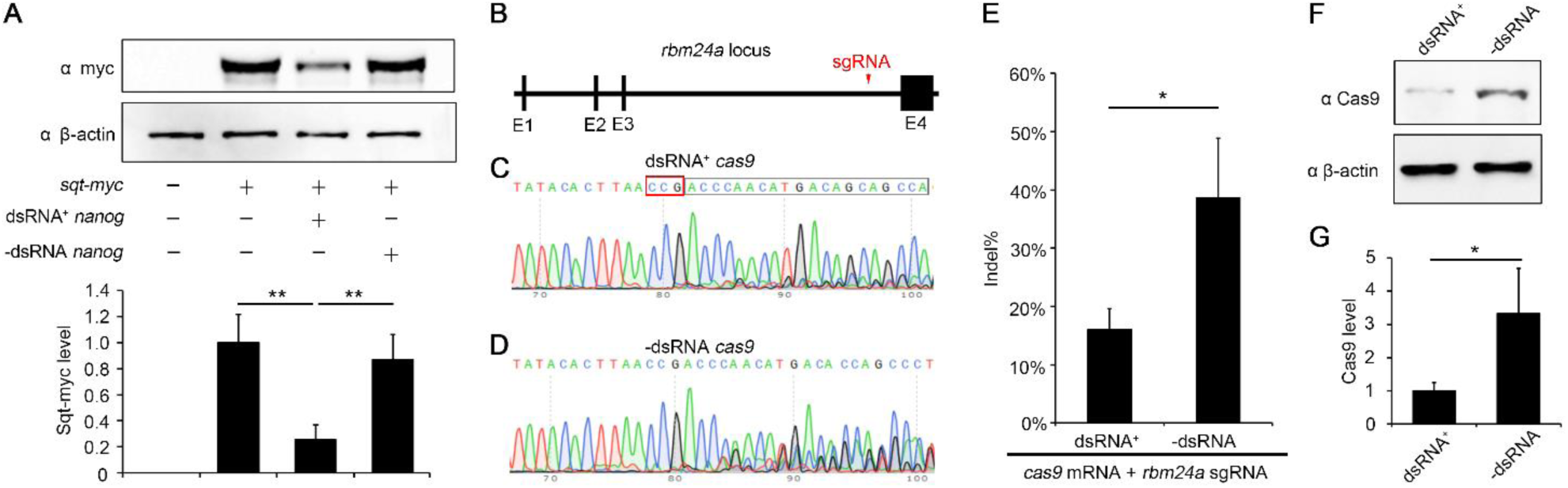
dsRNA in IVT mRNA reduces protein synthesis. (**A**) Co-injection of *squint-myc* mRNA along with dsRNA-contaminated *nanog* mRNA significantly diminishes Squint-myc protein production, while co-injecting dsRNA-free *nanog* mRNA does not have this effect. The upper panel presents a representative Western blot result, and the lower graph displays the quantification of protein levels. (**B**) Schematic representation of the *rbm24a* locus, highlighting the location of the sgRNA target site within the last intron. (**C** and **D**) Editing efficiency of dsRNA-positive and purified *cas9* mRNA co-injected with *rbm24a* sgRNA. (**E**) Quantification of editing efficiency indicates a significant increase following the removal of dsRNA. (**F** and **G**) Western blot analysis reveals that Cas9 protein production is markedly lower in the dsRNA-positive IVT mRNA group compared to the dsRNA-free group. Values are expressed as means ± SD. * p<0.05, ** p<0.01, Student’s *t*-test.

## Legends for Supplementary Videos

**Video S1. Live imaging of uninjected and *cas9* IVT mRNA-injected embryos.** We recorded the first ten cleavages and observed no discernible morphological changes, aside from a slight delay in the final few cell divisions of the *cas9* mRNA-injected embryo.

**Video S2. Live imaging of the gastrulation process of uninjected and *cas9* IVT mRNA-injected embryos.** During gastrulation, the lower-right embryo experienced an early burst of yolk cells, while the lower-left embryo had its yolk broken due to gastrulating movements. The upper-right embryos did not experience any bursting, but did exhibit epiboly delay and the emergence of large, swollen cells in the blastoderm during gastrulation.

**Video S3. PI staining of live embryos injected with dsRNA.** Time-lapse recordings were initiated at 6 hpf with the animal pole of the embryos oriented towards the objective lens. In a dsRNA-injected embryo, PI-stained red nuclei became visible over time, whereas they were absent in the uninjected control.

**Video S4. Isolated single dsRNA-stimulated cell underwent necrosis.** Single cells were isolated from both uninjected and dsRNA-injected embryos, and were observed under a Leica inverted microscope. The E3 medium contained Hoechst33342 and PI, with Hoechst33342 used to label the nucleus in live cells, while PI was used to label the nucleus only after cell membrane rupture.

## Supplementary Tables

**Table S1.**
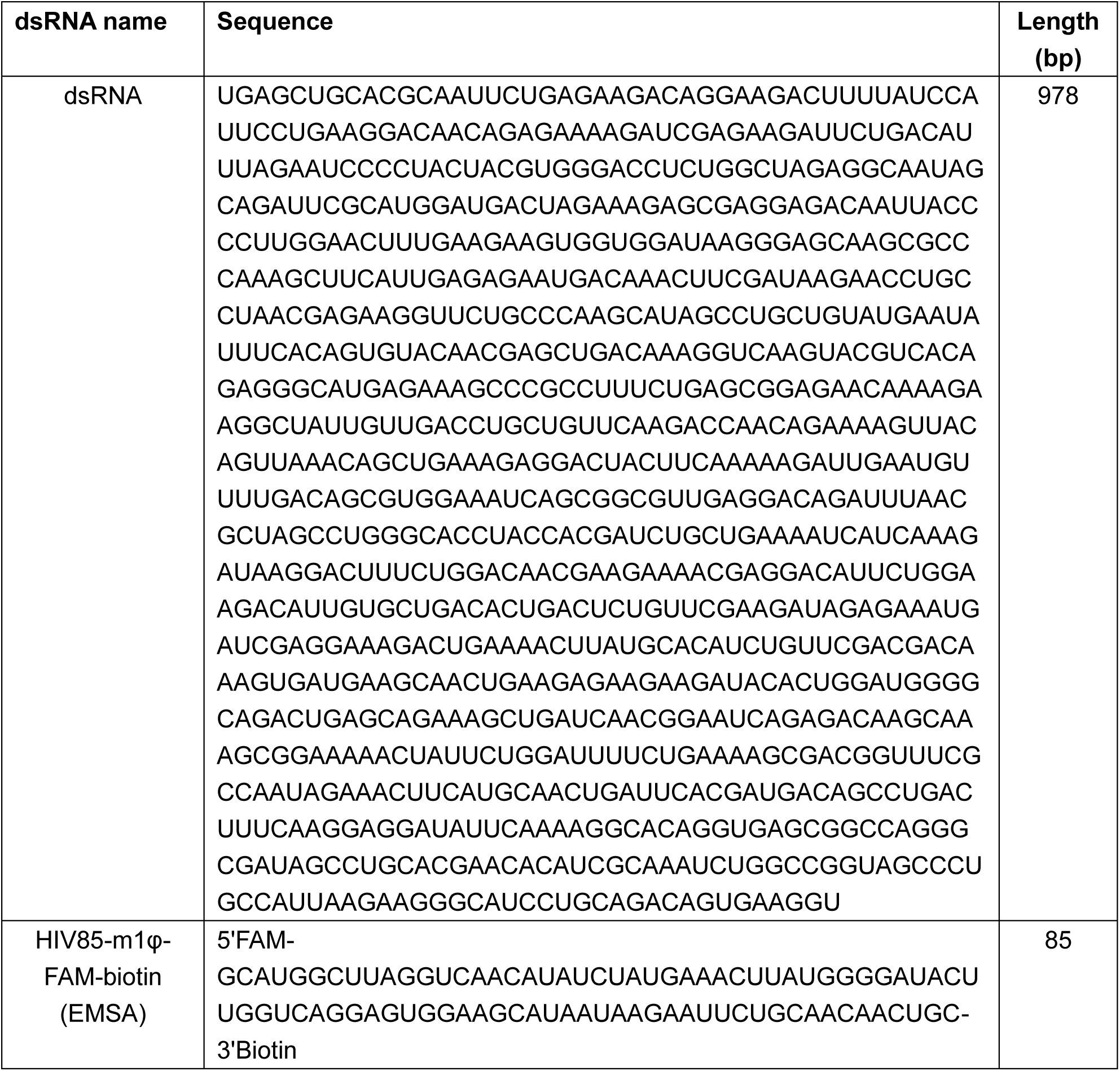
Sequences of dsRNAs.

**Table S2.**
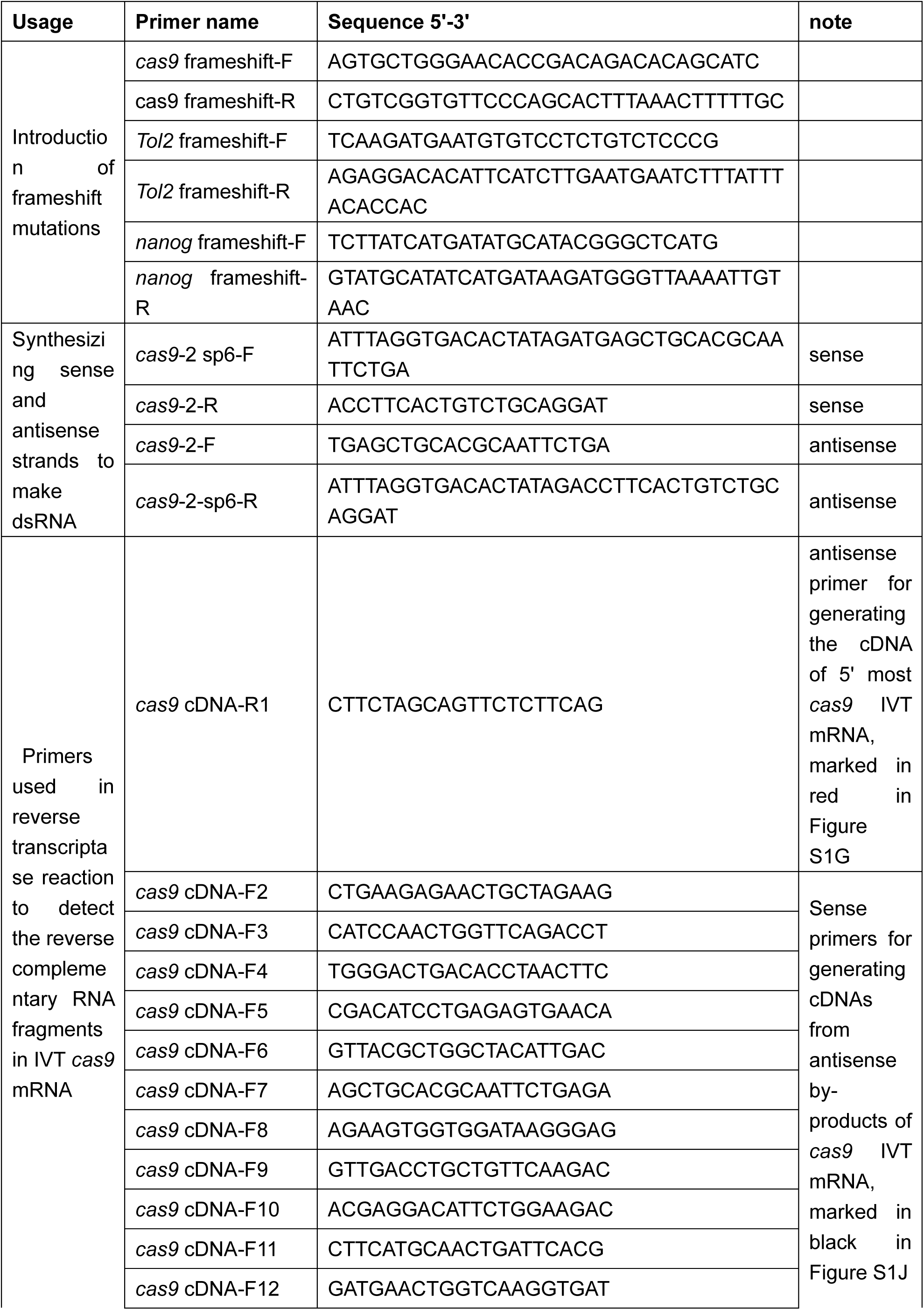
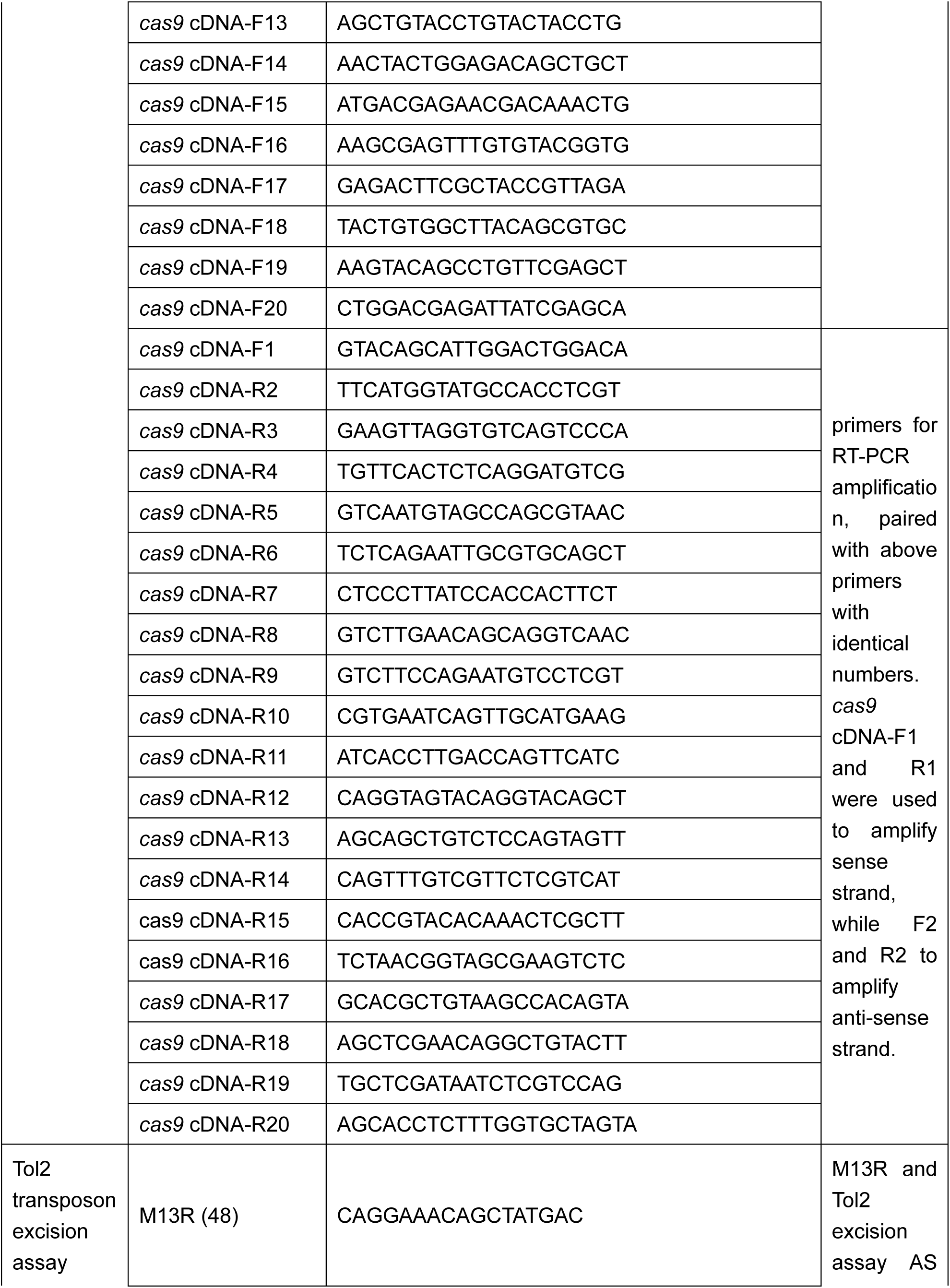
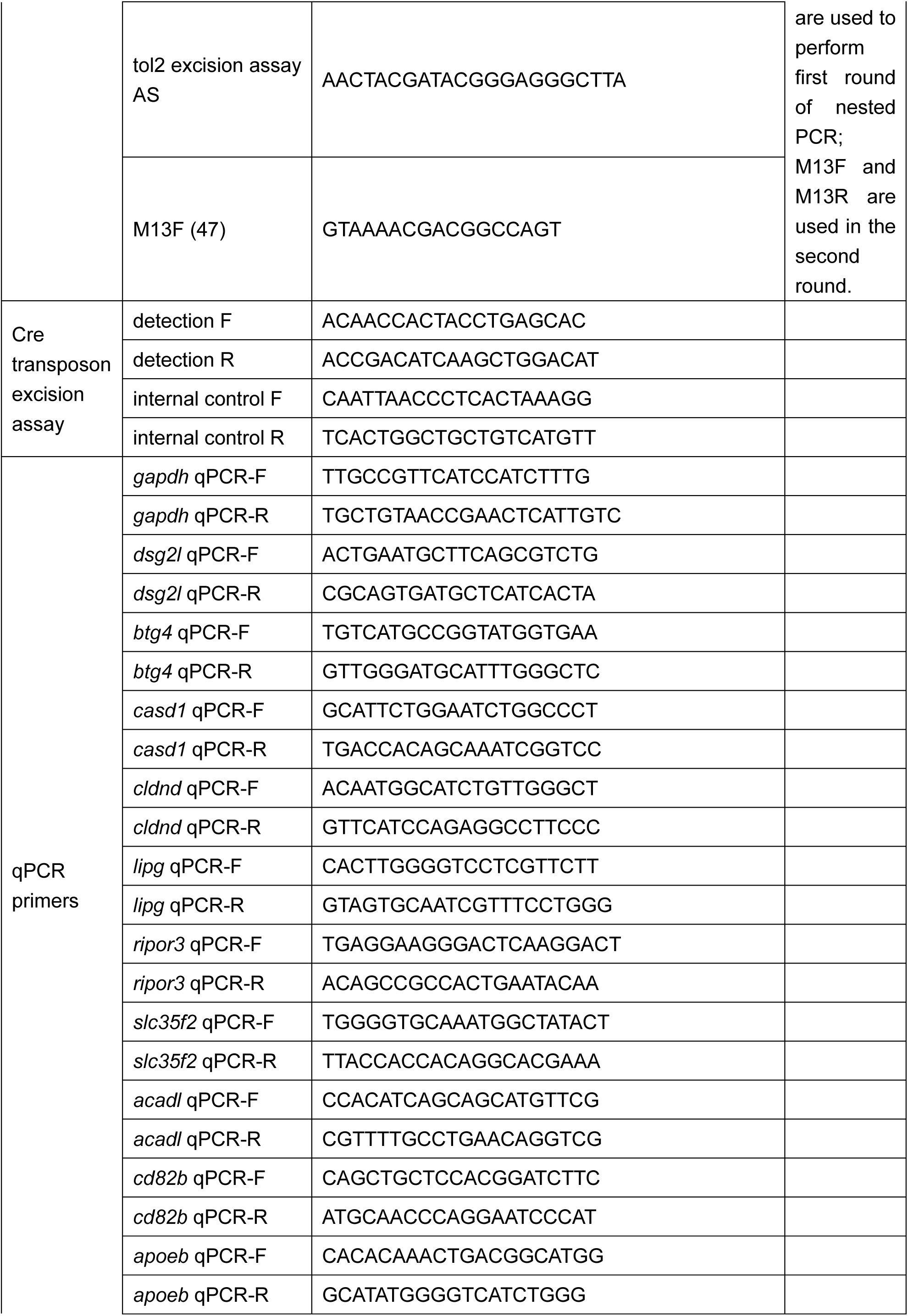
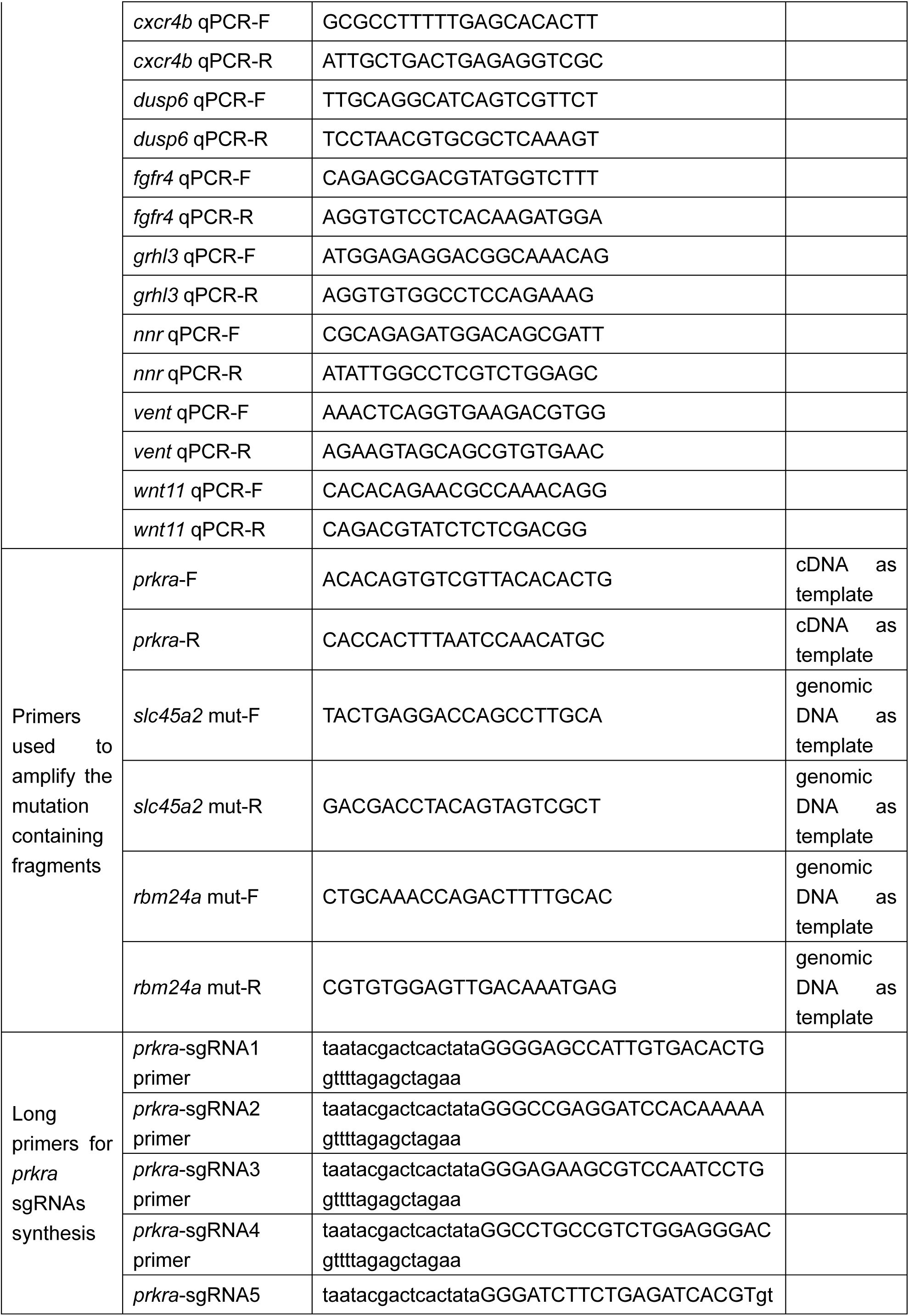
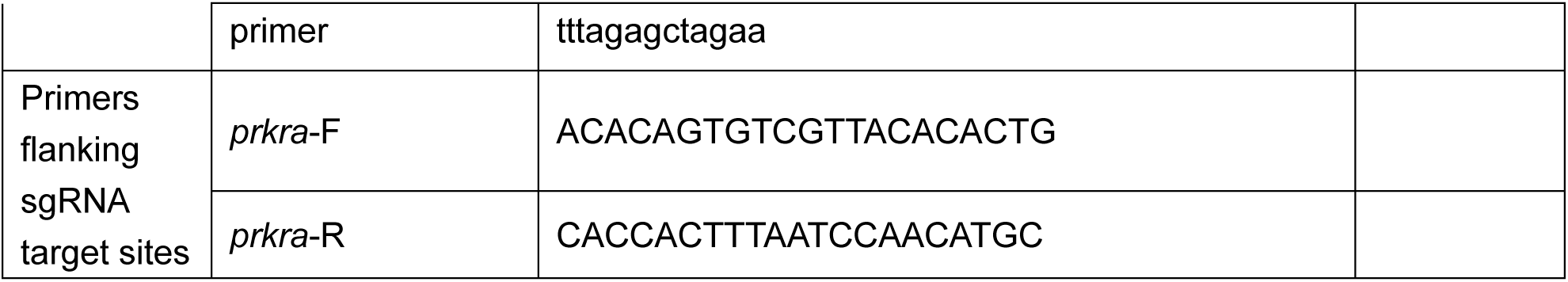
Primers used in this study.

## STAR Methods

### KEY RESOURCE TABLE

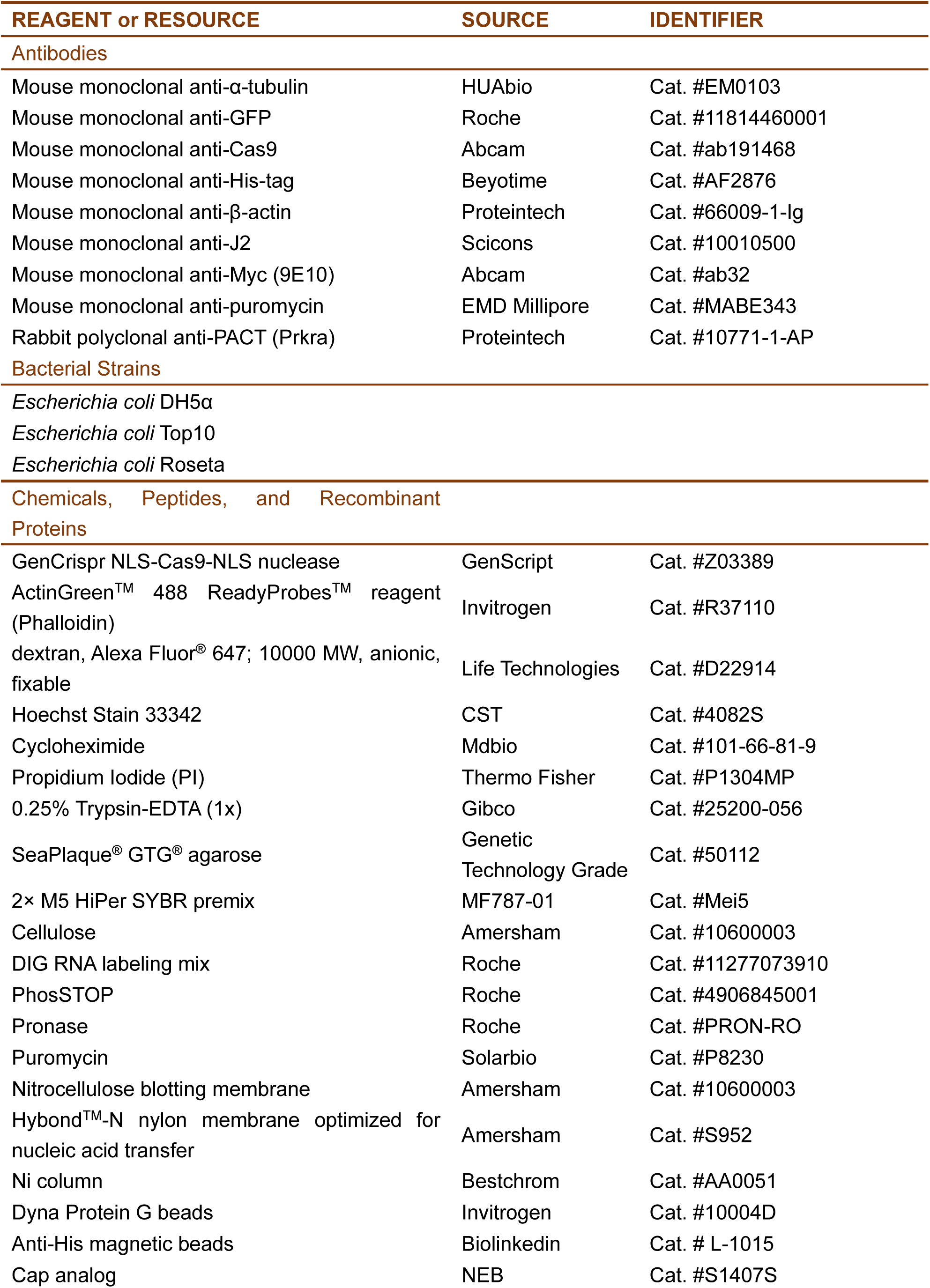
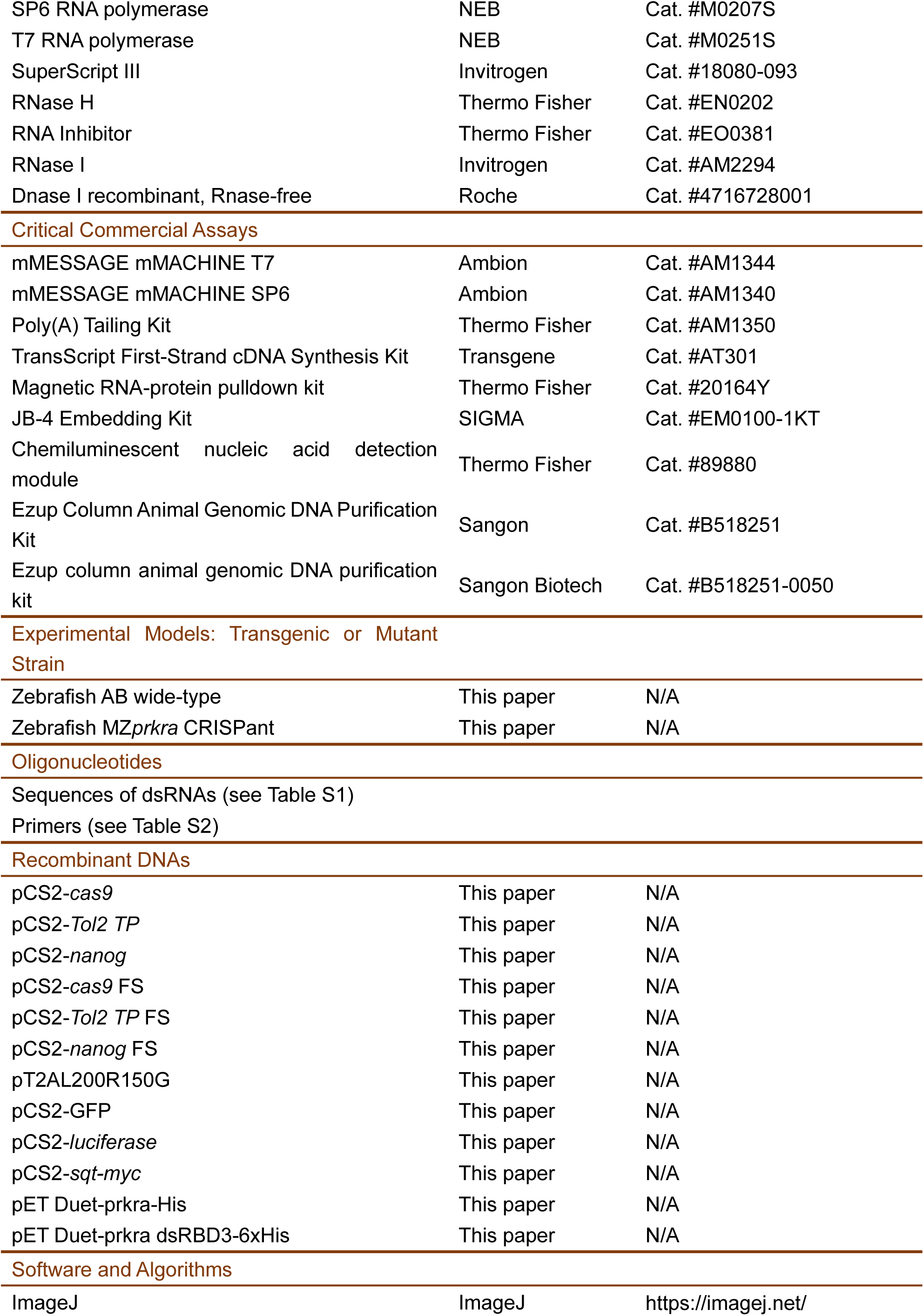
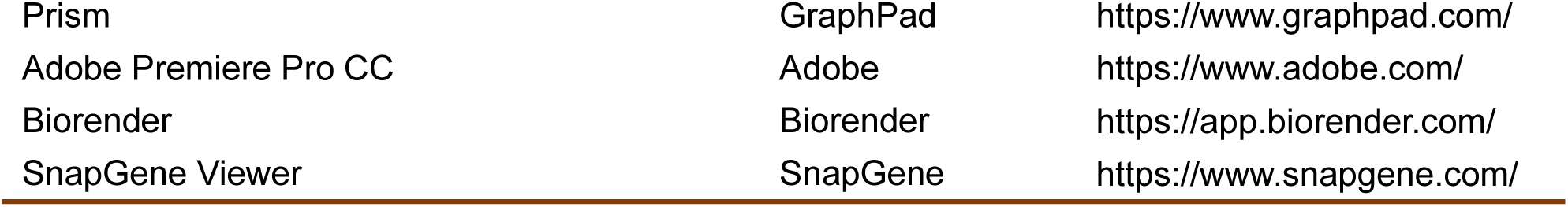

### RESOURCE AVAILABILITY

Further information and requests for reagents may be directed to and will be fulfilled by the lead contact Ming Shao.

#### Materials availability

All the constructs and reagents in this work are available from the lead contact upon request.

#### Data availablility

All data reported in this paper is available upon request to the lead contact.

### EXPERIMENTAL MODELS AND SUBJECT DETAILS

#### Animals

Zebrafish AB strain, as well as *nanog* and *prkra* mutant lines, were kept in standard breeding systems (Haisheng) at a temperature of 28 °C. All experiments adhered to the ARRIVE guidelines and were approved by the Ethics Committee for Animal Research of the School of Life Sciences at Shandong University (permit number SYDWLL-2023-069).

### METHOD DETAILS

#### Generation of maternal and zygotic mutants in zebrafish

To rapidly generate *prkra* maternal and zygotic mutants (MZ*prkra*), we utilized a CRISPant strategy. Five sgRNAs (Table S2) targeting the open reading frames (ORFs) of *prkra* were designed using the online software CRISPRscan (https://www.crisprscan.org/).^44^ These sgRNAs were synthesized via standard IVT and purification.44 The sgRNAs (100 pg/embryo in total) and Cas9 protein (200 pg/embryo) were injected into fertilized eggs. To assess the editing efficiency, injected embryos at 10 hpf were homogenized for genomic DNA preparation in 50 mM NaOH for 20 minutes at 95 °C, followed by neutralization with 1/10 volume of 1 M Tris-HCl (pH 7.5). Then, the DNA fragment flanking the sgRNA sites were amplified by PCR and subjected to Sanger sequencing. Next, the sequencing chromatograms were uploaded to http://shinyapps.datacurators.nl/tide/ to estimate the editing efficiency following the TIDE algorithm. The CRISPant F0 embryos were then raised to adulthood, and the resultant male and female fish were crossed to produce MZ*prkra* mutant embryos. These embryos were either raised to produce a stable mutant line or subjected directly to experimental assessment. They were genotyped by RT-PCR and Sanger sequencing to confirm the loss of Prkra function.

M*nanog* mutant embryos were generated following a previously established procedure.^31,45^ sgRNAs targeting *nanog* ORF were cloned into pU6-sgRNA vectors and then the sgRNA expression cassettes were excised and tandemly aligned into a pDest-3sgRNA vector via Golden Gate ligation. This vector was purified and coinjected with I-Sce I meganuclease for transgenesis. The GFP positive offspring were collected to examine the characteristic *nanog* mutant phenotype.

#### mRNA and dsRNA synthesis

Coding sequences of the genes used in this study were either PCR-amplified or directly synthesized, then subcloned into the pCS2 vector using a modified Gibson Ligation method.^46^ Recombinant vectors were linearized by Not I digestion, followed by purification with a PCR cleanup kit. The IVT reactions were performed with either the mMESSAGE mMACHINE SP6 or T7 kit (Cat. #AM1344 or Cat. #AM1340) according to the manufacturer’s instructions. Synthesized mRNAs were extracted with phenol-chloroform-isopentanol (25:24:1, v/v) followed by extraction with chloroform-isopentanol (24:1, v/v). The RNAs in the aqueous phase were then precipitated in 50% ice-cold isopropanol. After centrifugation at 12,000 g at 4 °C for 15 minutes, the RNA pellet was rinsed with 70% cold ethanol and resuspended in 15-20 μL of RNase-free water.

The m1φ-modified mRNAs were synthesized similarly as the UTP mRNAs except that rUTP was replaced by m1φ in the rNTP mix (10 mM rATP, 10 mM m1φ, 10 mM rCTP, 2 mM rGTP, 8 mM cap analog). Then the purified mRNAs were subjected to poly(A) tailing reaction following the procedure of the Poly(A) Tailing Kit (ThermoFisher, AM1350).

To generate dsRNAs, DNA templates were PCR-amplified from plasmids with T7 and SP6 promoter sequences incorporated into the 5’ or 3’ ends via primers. Sense and antisense RNAs were synthesized separately and then annealed at equal amount. The program for dsRNA annealing is as follow: 95 °C for 5 minutes, ramping down at 0.1 °C/s to 50 °C, and then quickly cooling to 4 °C.

#### dsRNA removal by cellulose chromatography

Fifty μg mRNAs were incubated with 50 mg of cellulose (Sigma-Aldrich, C6288) for 30 minutes in a buffer containing 10 mM HEPES (pH 7.2), 0.1 mM EDTA, 125 mM NaCl, and 16% (v/v) ethanol. The mixture was then centrifuged at 12,000 g at 4 °C, and the supernatant was collected. To precipitate RNAs from the supernatant, 1/10 volume of 4 M sodium acetate (pH 4.0) and 2.5 volume of absolute ethanol were added. After thorough vortexing, the mixture was incubated in a −20 °C refrigerator for 30 minutes, followed by centrifugation at 12,000 g for 15 minutes at 4 °C. The resulting precipitate was rinsed with 70% ethanol and dissolved in RNase-free water.

#### dsRNA detection

dsRNAs were detected following two strategies. First, they were detected by dot blot analysis using a dsRNA-specific J2 antibody (Scicons, 10010500). Briefly, equal amount of RNAs (200 ng) were spotted onto the Nylon membrane (Amersham, S952) using a 10 μL pipet tip and allowed to dry at room temperature. The membrane was then incubated with J2 antibody solution (1:2000, diluted in TBST) at 4 °C overnight, followed by washing with TBST three times every 10 minutes. The J2 antibody bound to the membrane was further detected by anti-mouse IgG-HRP diluted at 1:5000 in TBST. After rinsing the membrane every 10 minutes with TBST for three times, the dsRNA was visualized using ECL reagent (Meilunbio, MA0186) and imaged using a chemiluminescence imaging system (Sagecreation).

Second, dsRNAs were detected by amplifying the antisense strand of the *cas9* mRNA sequence using RT-PCR. Briefly, sense primers were designed to recognize the antisense RNA strands and were employed for cDNA synthesis. The amount of antisense *cas9* RNAs was quantified by qRT-PCR. In parallel, an antisense primer was utilized to synthesize cDNA specifically representing *cas9* mRNAs, which was subsequently quantified by qRT-PCR. The relative amount of antisense RNAs serves as an indicator of dsRNA levels, calculated as the ratio of antisense RNAs to mRNAs. The primer design and sequences are shown in Figure S1J and listed in Table S2.

#### Bright-field live-imaging

Embryos were dechorionated by pronase at a concentration of 1 mg/mL in Ringer’s solution for 10 minutes. They were then mounted in 1.5% low-melting agarose on a cover glass-bottomed petri dish, and their position was adjusted before agarose solidification. After adding 1/3x Ringer’s buffer, samples were loaded onto an inverse microscope (JiangNan, XD-202) for time-lapse recording. To acquire differential interference contrast (DIC) image, dechorionated embryos were mounted in 1.5% low-melting agarose on a concavity slide and photographed under a Leica upright microscope (Leica, DM2500) equipped with DIC apparatus. The videos were generated and further edited by Adobe Premiere Pro CC software.

#### Western blot

Embryos were dechorionated with their yolk cells manually removed. The resulting embryonic cells were collected in 1.5 mL centrifuge tubes and homogenized in a lysis buffer (100 mM NaCl, 0.5% NP- 40, 5 mM EDTA, and 10 mM Tris-HCl pH 7.5 with protease inhibitors). After clearing by centrifugation at 12,000 g for 10 minutes, cell lysates were boiled in 1x Laemmli buffer (10% SDS, 0.5% bromophenol blue, 50% glycerol, 5% β-mercaptoethanol, and 250 mM Tris-HCl, pH 6.8). Proteins were then separated by SDS-PAGE and transferred to a nitrocellulose membrane. The primary antibodies used for western blot can be found in the Key Resource Table.

#### qRT-PCR

Total RNA was extracted at the appropriate developmental stages, and cDNA synthesis was performed using the TransScript First-Strand cDNA Synthesis Kit (Transgene) according to the manufacturer’s instructions. qPCR was conducted with 2x M5 HiPer SYBR premix (MF787-01, Mei5) on a Q2000 real-time PCR system (LongGene). Primers used for gene expression analysis are listed in Table S2. Relative expression levels were calculated using the 2^-ΔΔCt^ method, with the expression of *gapdh* serving as the loading control.

#### Microinjection and cell transplantation

The injection needles were prepared using Narishigi needle puller (Puller PC-100). The melt-sealed end of the needle was clipped by tweezers, creating an open tip at a diameter of 5 μm. IVT mRNAs, dsRNAs or other reagents were loaded into the needle from the tip by the “Fill” function of the PLI-100A Picoliter Microinjector and then injected into 1-cell stage zebrafish embryos at a volumn of 1-2 nL per embryo.

Cell transplantation experiment was performed as described previously.^47^ Briefly, wild-type embryos were first labeled with Alexa647-dextran by microinjection at 1-cell stage. At high stage, cells in the blastodisc were aspirated and introduced into the marginal region of stage-matched unlabeled recipients. At 7 hpf, chimeric embryos were mounted in 1% low melting agarose on a cover slip for confocal imaging.

#### Tol2 transposon excision assay

100 pg of Tol2 TP IVT mRNA and 10 pg of pT2AL200R150G plasmid were co-injected into each 1- cell stage embryo. The injected embryos were raised to the shield stage and then lysed for DNA extraction using a genomic DNA extraction kit (Sangon, B518251). The extracted DNA was subsequently used as a template for nested PCR amplification. The primers used in this assay are listed in Table S1. M13R and Tol2 excision assay AS were employed for the first round of nested PCR, while M13F and M13R were used in the second round. The pre-amplification was set for 15 cycles. The PCR product from the pre-amplification was then diluted at 1:10 and used as a template for the second round of qPCR analysis. GAPDH was used as a loading control in the qPCR analysis.

#### Quantification of genome editing efficiency

The sgRNAs targeting the *slc45a2* open reading frame (ORF) and the third intron of *rbm24a* are listed in Table S1. DNA templates for the sgRNAs were synthesized by fill-in PCR following the guidelines on the CRISPRScan website. After IVT, purified sgRNAs were injected along with zebrafish codon-optimized *cas9* mRNA into the fertilized eggs. Embryos at 10 hpf were homogenized in 50 mM NaOH for 20 minutes at 95 °C, followed by the addition of 1/10 volume of 1 M Tris-HCl (pH 7.5). The DNA fragment flanking the sgRNA site was PCR amplified and sequenced. Sanger sequencing chromatogram files were submitted to http://shinyapps.datacurators.nl/tide/ to estimate the efficiency of the sgRNA.

#### Phalloidin, PI and Hoechst staining

For phalloidin staining, embryos were fixed with freshly prepared 4% paraformaldehyde and washed in PBST for three cycles of 10 minutes each. The embryos were then incubated with fluorescein-labeled (FITC) phalloidin for 30 minutes and washed again in PBST for three cycles of 10 minutes each. They were subsequently analyzed under a confocal microscope (Zeiss LSM880), and Z-stack projections were obtained using ZEN software.

For PI and Hoechst staining, live embryos, either uninjected or dsRNA-injected, were raised in E3 buffer containing 1 μg/mL PI and 1 μg/mL Hoechst33342. The embryos were subjected to fluorescence time-lapse imaging under an inverse microscope or mounted in 1.5% low-melting agarose and imaged using a Zess confocal microscope.

Embryos at 8 hpf were dissociated by trypsin-EDTA treatment (0.25% m/v) at 37 °C for 15 min with gentle pipetting. The dissociated cells from uninjected or dsRNA-injected embryos were washed and stained with 1 μg/mL PI and 1 μg/mL Hoechst33342 in E3 buffer. They were then subjected to time-lapse analysis using a Leica inverse microscope equipped with the Thunder function.

#### PI staining and quantification of cell necrosis

The number of PI-stained nuclei and the intensity of PI staining of each embryo can be used as cell necrosis indicators. 1 μg/mL of PI in E3 buffer was used to stain dsRNA-injected embryos until 7-8 hpf. The red fluorescent nuclei of each embryo were counted manually or the integrated optical density of the red fluorescence of each embryo was measured by Image J.

#### Electron microscopy

Embryos were fixed in 2.5% glutaraldehyde buffered in PBS for 1 to 4 hours at 4 °C. After washing in PBS five times for 20 minutes each, the embryos were further fixed in 1% osmium tetroxide. Following rinsing in PBS three times for 15 minutes each, the embryos were dehydrated through a graded ethanol series and subjected to two treatments with absolute acetone. They were then infiltrated with Epon 812 resin and polymerized using a temperature-rising program (12 hours at 36 °C, 12 hours at 50 °C, and 24 hours at 60 °C). The embedded samples were sectioned to a thickness of 100 nm and stained with 2% uranyl acetate and 3% lead citrate. Ultrathin sections were prepared using an ultramicrotome (PowerTome-XL) equipped with a glass knife and subsequently visualized under a focused ion beam scanning electron microscope (ZEISS, Crossbeam 550).

#### JB4 sectioning

Embryos were fixed overnight at 4 °C in PBS-buffered 4% paraformaldehyde. They were embedded in JB-4 resin following the manufacturer’s guide (Sigma, EM0100-1KT). 2 µm sections were obtained using an untramicrotome (PowerTome-XL) equipped with a glass knife and stained with hematoxylin and hydrophilic eosin Y.

#### ISH

In situ hybridization (ISH) was performed according to previously described protocols.^48^ Templates for synthesizing *prkra* probes were generated through PCR amplification. Digoxigenin-labeled probes were synthesized via IVT using the DIG-labeling mix (Roche) with SP6 RNA polymerase. Following treatment with DNase I to remove template DNA, the labeled probes were directly precipitated with 50% (v/v) isopropanol.

#### Puromycin incorporation assay

Embryos were exposed to 50 μg/mL puromycin from 2 to 4 hpf. Subsequently, they were collected after the removal of the yolk cell and lysed in protein immunoprecipitation (IP) buffer containing 10 mM Tris-HCl (pH 8), 150 mM NaCl, 1% NP-40, 1 mM EDTA, 10 μg/mL Aprotinin, 10 μg/mL Leupeptin, and 1 mM PMSF. Following denaturation at 95 °C for 5 minutes, samples were mixed with 1x Laemmli buffer (10% SDS, 0.5% bromophenol blue, 50% glycerol, 5% β-mercaptoethanol, and 250 mM Tris-HCl, pH 6.8). Puromycin-conjugated proteins were then visualized via western blot analysis using a puromycin-specific antibody (EMD Millipore, Cat. #MABE343).

#### Ribosome profile analysis

Sucrose was dissolved in polysome buffer (50 mM Tris-HCl, pH 7.0, 100 mM NaCl, 5 mM MgCl2, 100 μg/mL CHX) at concentrations of 10% and 60%. A sucrose density gradient was established using a 41Ti centrifuge tube on a Gradient Station (BIOCOMP) and allowed to reach equilibrium. Embryos at 5 hpf were harvested from both control and dsRNA-injected groups. Chorions were enzymatically digested using pronase, and the yolk cells were manually removed. The yolk-removed embryos were then exposed to 50 μg/mL CHX for 10 minutes before being homogenized on ice in lysis buffer containing 1% Triton X-100, 100 μg/mL CHX, 300 U/mL RNase inhibitor, 10 μg/mL aprotinin, 10 μg/mL leupeptin, and 1 mM PMSF. The lysate was then centrifuged at 12,000 g for 10 minutes at 4 °C. The resulting supernatant was carefully layered on top of the sucrose density gradient and subjected to ultracentrifugation at 240,000 g for 2 hours using an ultracentrifuge (Beckman, XPN-100). Subsequently, the centrifuge tube was transferred to an automated density fractionation system (BIOCOMP) for fraction separation, and the absorbance of each fraction was measured at 260 nm.

#### RNA pulldown

The detailed protocol is provided in the Pierce™ Magnetic RNA-Protein Pulldown Kit manual. Initially, 50 μL of streptavidin magnetic beads were transferred to RNase-free centrifuge tubes. These tubes were then placed on a magnetic separation stand for 30 seconds to facilitate separation, after which the supernatant was removed. The beads were washed twice with 20 mM Tris-HCl, pH 7.5 (50 μL for each wash). Subsequently, 1x RNA Capture Buffer was added to the beads, which were vortexed and incubated with 50 pmol of biotin-labeled RNAs under gentle rotation at room temperature for 20 minutes. After discarding the supernatant, the beads were washed twice again with 20 mM Tris-HCl, pH 7.5.

Concurrently, 400 de-yolked embryos were lysed in 400 μL of protein lysis buffer (10 mM Tris-HCl, pH 8, 150 mM NaCl, 1% NP-40, 1 mM EDTA, 10 μg/mL aprotinin, 10 μg/mL leupeptin, and 1 mM PMSF). The resulting lysate was centrifuged at 12,000 g for 10 minutes at 4 °C, and 130 μL of the cleared supernatant was diluted to a total volume of 400 μL in 1x protein-RNA binding buffer and 15% glycerol. This mixture was then incubated with the beads at 4 °C under rotation for 1 hour. Following the incubation, the beads were washed twice with 100 μL of 1x washing buffer with gentle mixing. Finally, the beads were incubated with 50 μL of elution buffer at 37 °C for 20 minutes. The eluate was subsequently analyzed by western blot.

#### Recombinant protein expression and purification

The pET-Duet plasmid was linearized using Kpn I and Nco I restriction enzymes. A 6x His tag sequence was incorporated at the 3’ end of the coding regions for *prkra*. The Rosetta strain of E. coli, transformed with the plasmids, was cultured overnight at 37 °C in LB medium and then inoculated into 1 L of fresh LB for amplification until an optical density (OD) of 0.6 was reached. To induce protein expression, IPTG (final concentration of 1 mM) was introduced into the culture and incubated for an additional 26 hours at 20 °C with agitation. The bacteria were harvested by centrifugation at 2,800 g for 10 minutes at 4 °C, and the pellets were resuspended in 10% glycerol before undergoing a second centrifugation at 1,850 g for 20 minutes at 4 °C. The cell pellets were weighed and subsequently resuspended in AP buffer (50 mM Tris-HCl, pH 7.2, 400 mM NaCl, 40 mM imidazole, 10% glycerol, and 0.02% NP-40). The cells were homogenized using lysozyme solution (1 mM DTT, 1 mM PMSF, and 1 mg/mL lysozyme in AP buffer), followed by sonication with an ultrasonic homogenizer (JY92-IIN) set to a cycle of 5 seconds ON and 10 seconds OFF for a total of 4 minutes at a power setting of 20%. After centrifugation at 24,000 g for 1 hour at 4 °C, the supernatant was filtered through a 0.45 μm membrane. In parallel, a Ni column was prepared, washed twice with ddH2O, and then once with AP buffer. The cleared supernatant was added to the column and incubated at 4 °C for 1 hour to facilitate protein binding. The column was drained and washed twice with buffer A (1 mM DTT and 1 mM PMSF in AP buffer), followed by the addition of cold buffer B (2 mM ATP-Na, 1 mM MgAc, 1 mM DTT, and 1 mM PMSF in AP buffer) and incubation at 4 °C for 10 minutes to activate the protein. Following two washes with buffer C (1 mM DTT in AP buffer), proteins were eluted slowly using the elution buffer (50 mM Tris-HCl, pH 7.2, 400 mM NaCl, 400 mM imidazole, 10% glycerol, and 0.02% NP-40). For long-term storage, glycerol was added to the protein solution to achieve a final concentration of 50%, and the samples were stored at −80 °C.

#### EMSA assay

An 85 bp dsRNA sequence was derived from the Vpr gene of HIV-1, with the sequence: GCAUGGCUUAGGUCAACAUAUCUAUGAAACUUAUGGGGAUACUUGGUCAGGAGUGGAAGCAUAAUAAGAAUUCUGCAACAACUGC. Two forward ssRNA strands were chemically synthesized by Sangon, either with or without m1φ modification. Both synthesized ssRNAs were labeled with FAM at the 5’ end and with biotin at the 3’ end. The reverse ssRNA strands were in vitro transcribed with or without m1φ modification. The m1φ modified and unmodified forward and reverse ssRNAs were then annealed to produce fluorophore-labeled m1φ dsRNAs or UTP dsRNAs. The dsRNAs and Prkra protein were mixed in 1x binding buffer (10x storage: 80% glycerol, 100 mM MgCl2, 5 M NaCl, 0.5 M EDTA, 1 M DTT, 1 M Tris-HCl, pH 8.0) and incubated at room temperature for 15 minutes. Subsequently, the protein-RNA mixture was electrophoresed using a 6% native PAGE gel at 80 V for 2 hours in 1x running buffer (25 mM Tris, 0.5 M glycine). Finally, gel imaging was performed using a full-spectrum gel imager (Invitrogen, iBright 1500).

#### Quantification and Statistical Analysis

All experiments were performed at least in duplicate with similar results. Data plotting and statistical tests were performed using GraphPad Prism 9. Statistical information is described in each figure legend.

