## Supplementary material for "N1-methylpseudouridine mRNA modification enhances efficiency and specificity of gene overexpression by preventing Prkra-mediated global translation repression": Uncropped gels and blots

**Uncropped blots for Figure 1F**

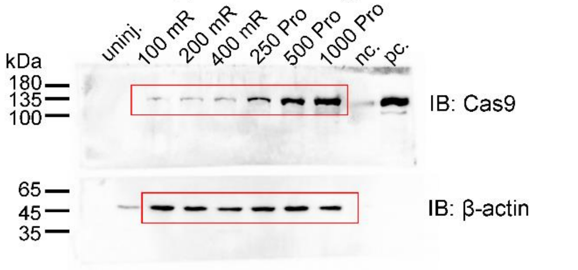


**Uncropped blots for Figure 1H**

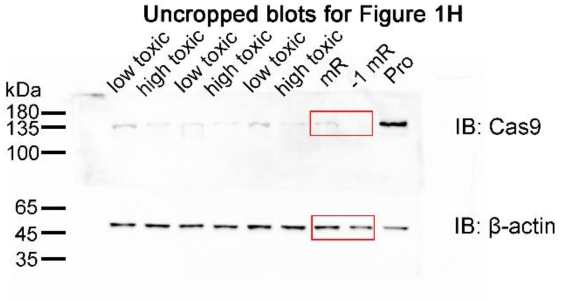


**Uncropped blots for Figure 1J**

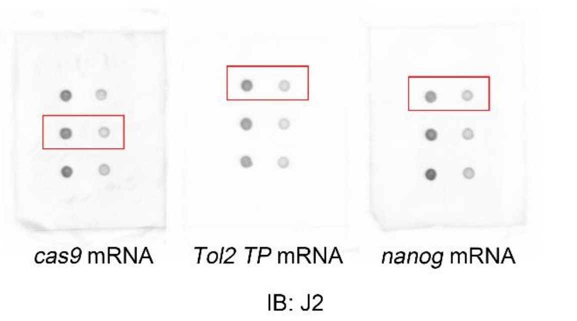


**Uncropped blots for Figure 3E**

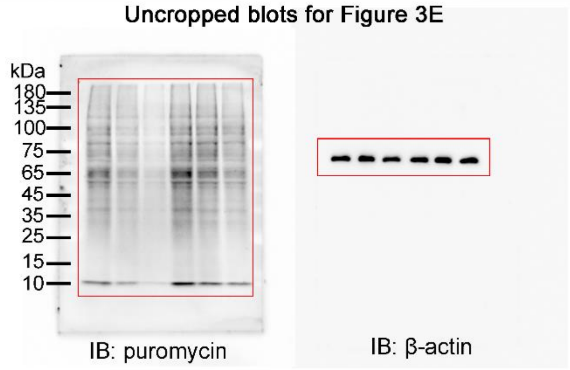


**Uncropped blots for Figure 5A**

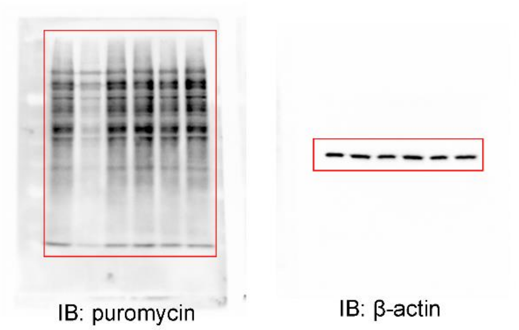


**Uncropped blots for Figure 5E**

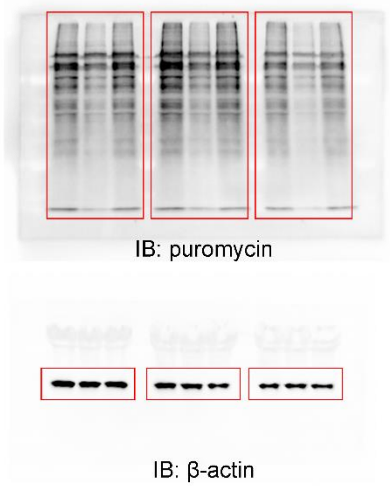


**Uncropped blots for Figure 6A**

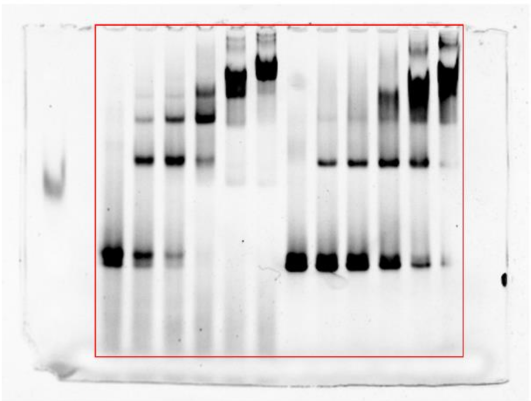

**Uncropped blots for Figure 6C**

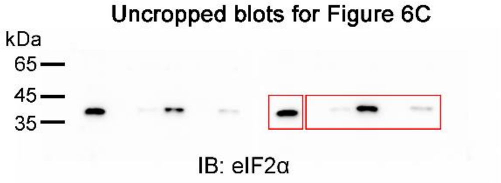


**Uncropped blots for Fig. 6D**

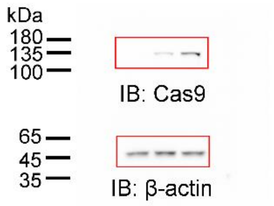

### Uncropped blots for Figure 6E

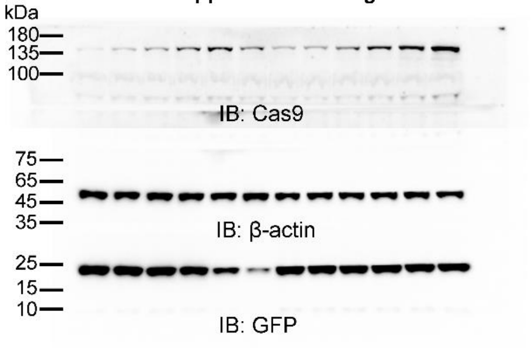


### Uncropped blots for Figure S1E

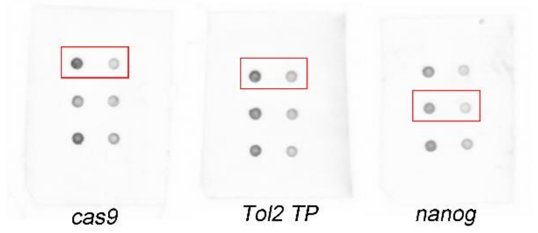


### Uncropped blots for Figure S1H

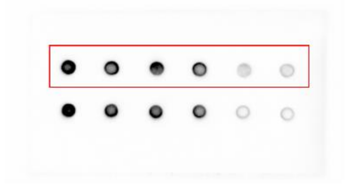

### Uncropped blots for Figure S6A

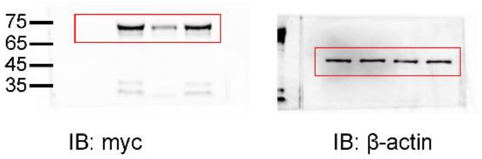


### Uncropped blots for Figure S6F

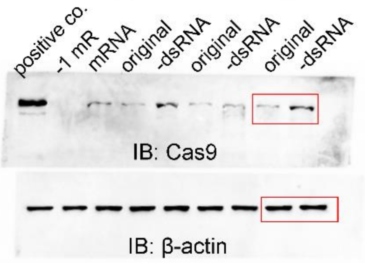


Same blot as Fig 1H, horizontally flipped

### Uncropped blots for Figure S1K

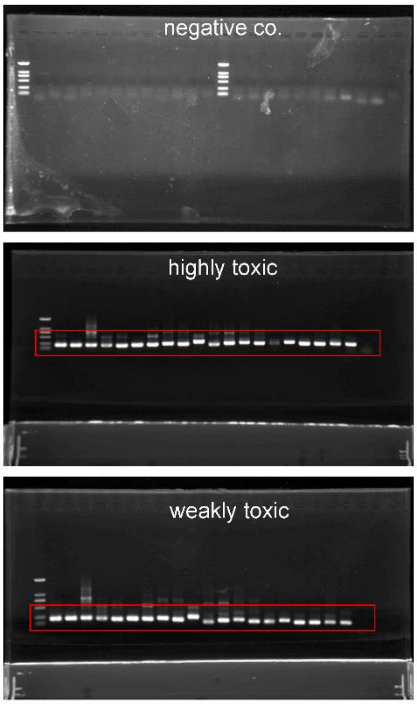
